# Systematic image perturbations reveal persistent gaps between human and machine vision

**DOI:** 10.64898/2026.08.01.742229

**Authors:** Mugihiko Kato, Biyu J. He

**Affiliations:** Department of Neuroscience, New York University Grossman School of Medicine, New York, NY 10016, USA; Department of Neurology, New York University Grossman School of Medicine, New York, NY 10016, USA; Department of Radiology, New York University Grossman School of Medicine, New York, NY 10016, USA; Department of Biomedical Engineering, NYU Tandon School of Engineering, New York, NY 11201, USA

## Abstract

Deep neural networks (DNNs) are promising computational models for understanding visual object recognition. Yet, whether DNNs use similar visual cues for object recognition as humans do remains unknown. We created an image set that systematically untangles global shape, internal parts, and texture information, and compared human recognition behavior against >200 DNNs spanning diverse architectures, training diets, and training objectives. No DNNs replicated humans’ cue-reliance profile, including those with recurrence or specialized training. Fine-tuned text-image contrastive-trained models, regardless of architecture, were most human-like overall, but lost their human-alignment when the global shape was disrupted. Strikingly, all DNNs substantially underperformed humans when the global shape cue alone was critical to object recognition. Furthermore, alignment with ventral stream neural recordings in an existing database did not predict alignment to human behavior, and model performance does not always predict its human-alignment. Together, these findings reveal systematic and persistent differences between human and machine vision.

## INTRODUCTION

Deep neural networks (DNNs) have become the leading computational model for primate vision^1^. They rival human observers in object recognition accuracy on certain benchmark tasks^2–4^ and exhibit hierarchical representations reminiscent of the primate ventral visual stream^5–10^. These properties make DNNs crucial models for investigating the computational principles underlying biological vision.

However, how DNNs align with or deviate from biological vision is currently unclear. While earlier studies showed that unlike humans, DNNs often fail under modest image perturbations, such as added noise, style alterations, or other minor distortions^11–16^, whether these pitfalls continue to apply to state-of-the-art DNNs is currently unknown. This is an urgent question given that model architectures and training approaches are evolving rapidly and many of these earlier studies focused on standard feedforward convolutional neural networks (CNNs) alone. Germane to this topic is the question of whether current DNNs use the same diagnostic cues for image recognition as humans do, which has implications on whether DNNs capture the core computational mechanisms of human visual object recognition.

Importantly, recent work showed that neither improving image classification accuracy alone nor scaling model size or training dataset size is sufficient to produce full alignment with human recognition behavior or with the primate ventral visual stream^1,11,17–19^. This suggests that DNNs might use systematically different diagnostic cues for image recognition as compared to humans. Understanding the potential discrepancies here can provide clues for developing more human-aligned machine vision, which can in turn serve as better models of human visual perception in both health and diseases.

Previous works comparing the performance of standard ImageNet-trained feedforward CNNs with humans have exposed two main discrepancies in visual-cue usage: First, when presented with stimuli containing conflicting shape and texture information, CNNs prioritize texture whereas human observers show a strong shape bias^20,21^. Consistent with this, CNNs’ outputs and hidden-layer activations are relatively insensitive to spatial scrambling^22–24^. Second, when presented with silhouettes or geometric shapes, CNNs are highly vulnerable to local contour disruptions that preserve the global shape, whereas humans are more sensitive to disruptions in the global shape^21,25–28^. These results demonstrate clear mismatches in the cue reliance profile between humans and standard feedforward CNNs, and performance on some of these tasks has become a benchmark for evaluating how human-like DNNs are^11^.

However, there are important gaps in the evaluation approaches used in previous studies. First, the widely used shape-texture conflict stimuli were constructed using only texture images that pretrained CNNs could classify correctly^20^, ensuring that the texture information in these stimuli is easily accessible by the CNNs. In addition, since the two visual cues are directly pitted against each other, models that perform well on this benchmark (i.e., having a high shape bias) might do so because of an inability to leverage the texture information instead of being genuinely able to utilize the shape information. To address this issue, shape and texture cues must be independently manipulated. Second, studies demonstrating feedforward CNNs’ bias toward local shape over global shape information often relied on a small number of object classes or image examples^21,25,27^. Given that diagnostic features vary widely across viewpoints, styles, and object classes, an unbiased evaluation using systematic perturbations of multiple visual cues across a number of image classes is necessary to robustly compare DNNs’ and humans’ object recognition behavior. Finally, as previously pointed out, it is important to evaluate image-wise error patterns instead of the overall accuracy alone^11,29^: a model that disagrees with humans on which images are easy or hard is not human-like, even if it has a similar overall accuracy to humans.

In addition to gaps in the evaluation approach, previous studies often focused on the evaluation of standard feedforward CNNs. Crucially, a wide range of models have been developed recently, which vary in architecture, training objective, and training data. For instance, models trained jointly on images and natural language (e.g., Contrastive Language-Image Pre-training; CLIP) narrow the performance gap with humans under certain image perturbations^11,28^ and better predict cortical visual responses^10,30,31^. In addition, training on noise-added, blurred or stylized images enhances the model’s shape bias as assessed by shape-texture cue-conflict images^20,29,32^. Finally, biologically inspired recurrent connections have been added to feedforward CNNs, improving performance in certain conditions and enhancing alignment with neural data recorded from the ventral stream^33,34^. However, to date, a comprehensive comparison of different DNN models against humans in their object recognition behavior using systematic manipulations of different visual cues is missing. Such an investigation would reveal whether DNNs use similar visual cues for object recognition as humans do, and which model components (e.g., architecture/training diet/objectives) affect human-likeness in visual cue usage and object recognition behavior.

To fill in these gaps, we created a large, unbiased image set spanning a diverse range of object classes and applied controlled perturbations to disrupt texture, global shape (henceforth “**shape**”), and local shape (henceforth “internal **parts**”) information either individually or in pairs. Under these manipulations, we compared image-wise error patterns and the overall recognition accuracy between human observers and >200 DNNs encompassing a wide variety of architectures, training objectives and training data. In a second experiment, we created manipulated images that fully decouple local and global shape cues.

Using this tailored image set, we reveal both universal failures of current DNNs and shortcomings specific to individual DNN classes, including models often regarded as well-aligned with biological vision. These findings highlight new directions for developing models that more closely match human object recognition behavior.

## RESULTS

### Overview of Methods and Experiment 1

Experiment 1 tested how disrupting shape, texture, and internal parts affects object recognition in humans and DNNs. First, we curated an image set spanning 48 basic object categories covering animate (both mammals and non-mammals) and inanimate (both manmade and natural) domains (**Fig. S1A**), with 5 exemplar images per basic category (240 exemplar images in total). From each exemplar image, we created six perturbation conditions, each disrupting one (**Fil**, **Lin**, **SilTex**) or two (**Sil**, **Tex**, **LinFil**) of three crucial visual cues: global shape (**S**), texture (**T**), and local shape (/internal parts, **P**) (**Figs. 1A, S2**). Specifically, **Fil** (filled), **Lin** (line-drawing), and **SilTex** (silhouette-texture) images disrupt global shape, texture, and internal parts (i.e., local shape), respectively. **Sil** (silhouette), **Tex** (texture), and **LinFil** (line-filled) images preserve only global shape, texture, and internal parts information, respectively. Detailed definitions of each visual cue and the procedures used to disrupt them are described in Methods, *Image manipulation* section, and **Fig. S1B**.

**Fig. 1.**
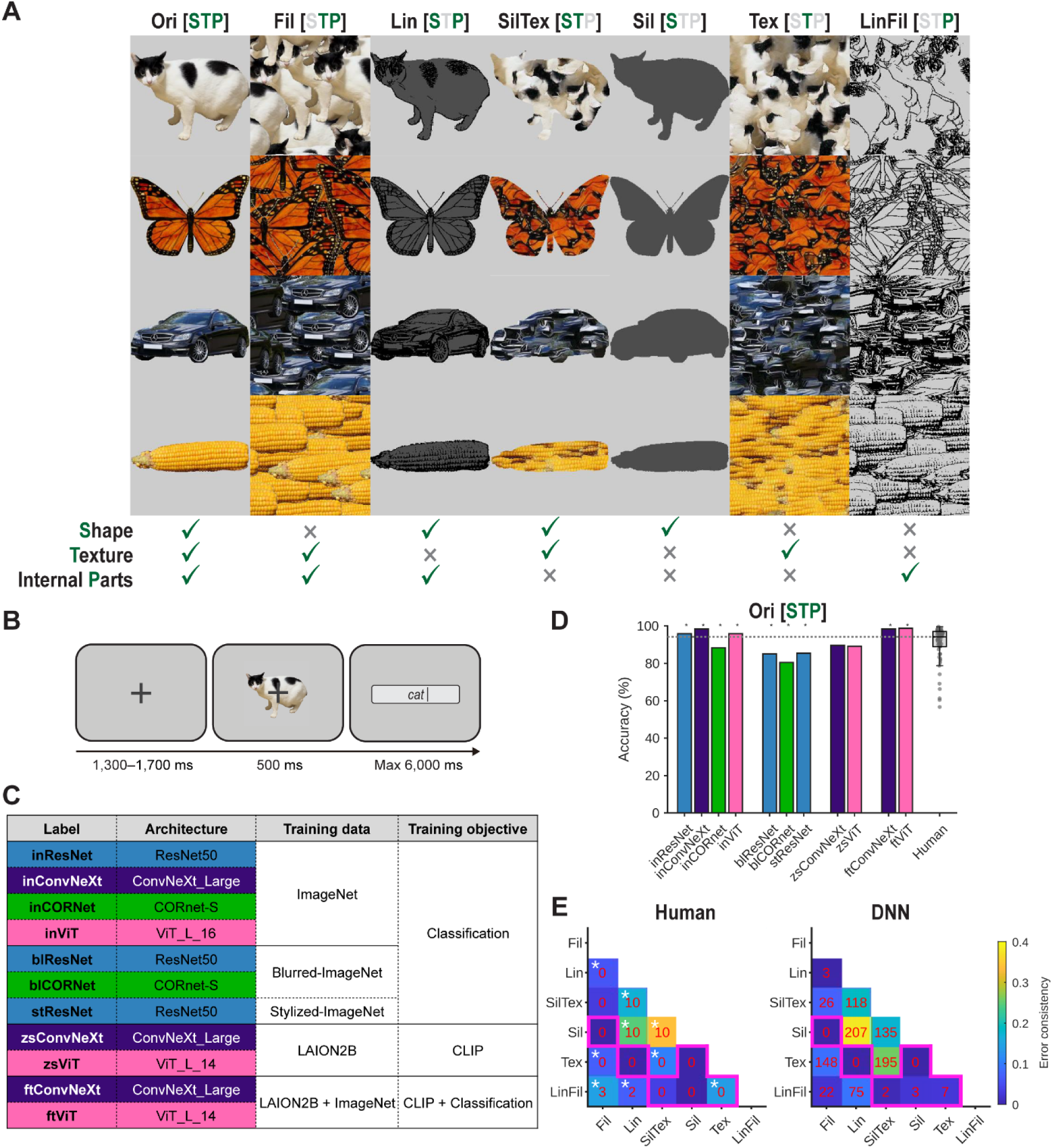
Overview of Experiment 1, accuracy in the original condition, and validation of image manipulations. **(A)** Seven image-manipulation conditions used in Experiment 1. For each condition, green letters adjacent to the condition label (S: shape; T: texture; P: internal parts) and green check marks beneath the example image indicate which feature(s) are preserved. See Methods and Fig. S1B for image-generation details. Note that the examples shown here are for illustrative purposes only and do not come from the actual stimulus set, owing to copyright restrictions. Additional examples are provided in Fig. S2. **(B)** Trial timeline of the object-recognition task: after a fixation delay, subjects viewed an object image for 500 ms and then typed the object name that first came to mind. **(C)** List of the 11 DNNs primarily examined. See Methods for details. **(D)** Recognition accuracy of the 11 DNNs and human observers in the original (Ori) condition. In all box plots, the center line denotes the group median; the box limits denote the interquartile range; and the whiskers extend to the maximum and minimum data points within 1.5 times the interquartile range. The overlaid dots represent individual subjects (N = 60). The horizontal dashed line indicates human median accuracy. Stars above the bars denote significant differences between DNN accuracy and human accuracy (two-sided Wilcoxon signed-rank tests; p < 0.05, Bonferroni-corrected; exact statistics in the supplementary Excel file). **(E)** Error consistency averaged across pooled human subjects (left) and across all 209 DNNs (right). The overlaid red numbers indicate the number of observers showing significant error consistency (humans, out of 10; DNNs, out of 209; permutation tests; p < 0.05; FDR-corrected; see Methods for details; exact statistics in the supplementary Excel file). For human data (left), white asterisks indicate significant error consistency based on a group-level statistical test (Wilcoxon signed-rank test; p < 0.05; FDR-corrected). The pink outline indicates condition pairs for which no visual cues are expected to be shared.

Human recognition behavior was measured in an online experiment (**Fig. 1B**). Each of the 60 participants completed ∼1 hr of the main task, during which she/he viewed all 240 exemplar images, with each image assigned to one of the six manipulated conditions, followed by all 240 images shown in the original (**Ori**) condition (**Fig. S1C**). To compare with DNNs, data from six subjects who together completed the full set of 240 × 6 manipulated images (i.e., including each exemplar image in all 6 manipulation conditions) were pooled into a single “pooled subject” (see Methods). Importantly, different “pooled subjects” consist of non-overlapping sets of subjects and therefore are statistically independent. This design was necessary to avoid potential learning effects across repeated presentations of the same exemplar image in different manipulation conditions in human participants.

Unlike human subjects, DNNs do not learn at test time, so the same DNN can be tested on the entire image set without interference between test conditions. Thus, each DNN was tested on the entire image set (240 exemplars × 7 conditions), and their top-1 predictions were recorded. To fairly compare human and DNN behavior, their answers were evaluated based on a lexical hierarchy defined by WordNet^35^ (see Method for details). We first focused our evaluation on eleven representative DNNs (**Fig. 1C**), which varied in both architecture (**Fig. 1C**, 2^nd^ column) and training data/objective (**Fig. 1C**, 3^rd^ and 4^th^ columns). Model architectures included feedforward CNNs (ResNet, ConvNeXt), a recurrent CNN (CORNet), and a Vision Transformer (ViT_L). Models were trained using either a classification task [based on standard ImageNet (labeled here as “in”), blurred images (“bl”), or stylized images (“st”)], contrastive language-image pretraining (CLIP) alone (labeled as ‘zs’ for zero-shot), or CLIP training fine-tuned with classification task training (labeled as ‘ft’ for fine-tuned). Models are denoted by the abbreviated labels in **Fig. 1C**, 1^st^ column, and the rationale for their selection is explained below. Afterward, we also conducted a larger-scale analysis that included a wider range of 209 DNNs to confirm the generalizability of our findings (Table S1).

In the **Ori** condition, recognition accuracy was high for humans and all models, although several models performed slightly but significantly below (inCORnet, blResNet, blCORnet, stResNet) or above (inResNet, inConvNeXt, inViT, ftConvNeXt, ftViT) the human accuracy level (**Fig. 1D**; p < 0.05; two-sided Wilcoxon signed-rank test, Bonferroni-corrected; exact statistics in the supplementary Excel file). To assess robustness to perturbations independently of baseline performance, for each human/DNN observer, we excluded any images misclassified in the **Ori** condition from subsequent analyses.

### Validation of image manipulations

Prior to evaluating humans vs. DNNs on our image set, we performed a validation analysis of our image manipulation procedure. A potential concern is that the disruption of a particular visual cue may be incomplete, in which case the residual information related to that cue could still drive recognition behavior. To examine whether this might be the case, we computed image-wise error consistency for every pair of image conditions, separately for each pooled human subject and for each of the 209 DNNs assessed in this study. Between-condition, within-subject error consistency measures the extent to which an observer makes errors on the same exemplar image across two conditions^11,36^.

We reasoned that if two image conditions preserve distinct diagnostic cues, successful identification of an exemplar image in one condition should not predict its identification in the other condition; accordingly, between-condition, within-subject error consistency should be near chance. Thus, if our image manipulations were successful, error consistency for condition pairs that share no preserved cue (i.e., **Fil-Sil**, **Lin-Tex**, **SilTex-LinFil**, **Sil-Tex**, **Tex-LinFil**, and **Sil-LinFil**) should be close to the chance level. To test this, for each pair of image conditions and each human/DNN observer, we computed between-condition error consistency and assessed its statistical significance by comparing with a null distribution obtained by permutation (for details, see Methods). None of the 10 pooled human subjects showed significant error consistency for any of these six condition pairs (**Fig. 1E**, left, pink outlines). Similarly, only a very small number of the 209 DNNs showed significant error consistency for these condition pairs (0–7 out of 209; **Fig. 1E, right**, pink outlines). Thus, for condition pairs that are supposed to share no diagnostic cues, distinct visual information appears to drive image recognition such that their error consistency is near chance, confirming that our image manipulations disrupted the visual cues as intended.

For human data, we additionally performed a group-level Wilcoxon signed-rank test on error consistency across the 10 pooled subjects. Significant between-condition error consistency was found in 9 condition pairs, with 8 of them being condition pairs with shared visual cues (**Fig. 1E**, left, white asterisks). Interestingly, there was significant error consistency in the **Tex**-**LinFil** condition pair (mean error consistency = 0.142), which is not supposed to have any shared visual cues, suggesting potential imperfection in cue dissociation between these two conditions. However, this between-condition, within-subject error consistency was much lower than the within-condition, between-subject error consistency observed in either the **Tex** or the **LinFil** condition (mean = 0.329 and 0.439; **Fig. 2F and G**, respectively; see the next section). This suggests that, although some overlap in error patterns exists between **Tex** and **LinFil** conditions, our dissociation of visual cues was largely successful.

**Fig. 2.**
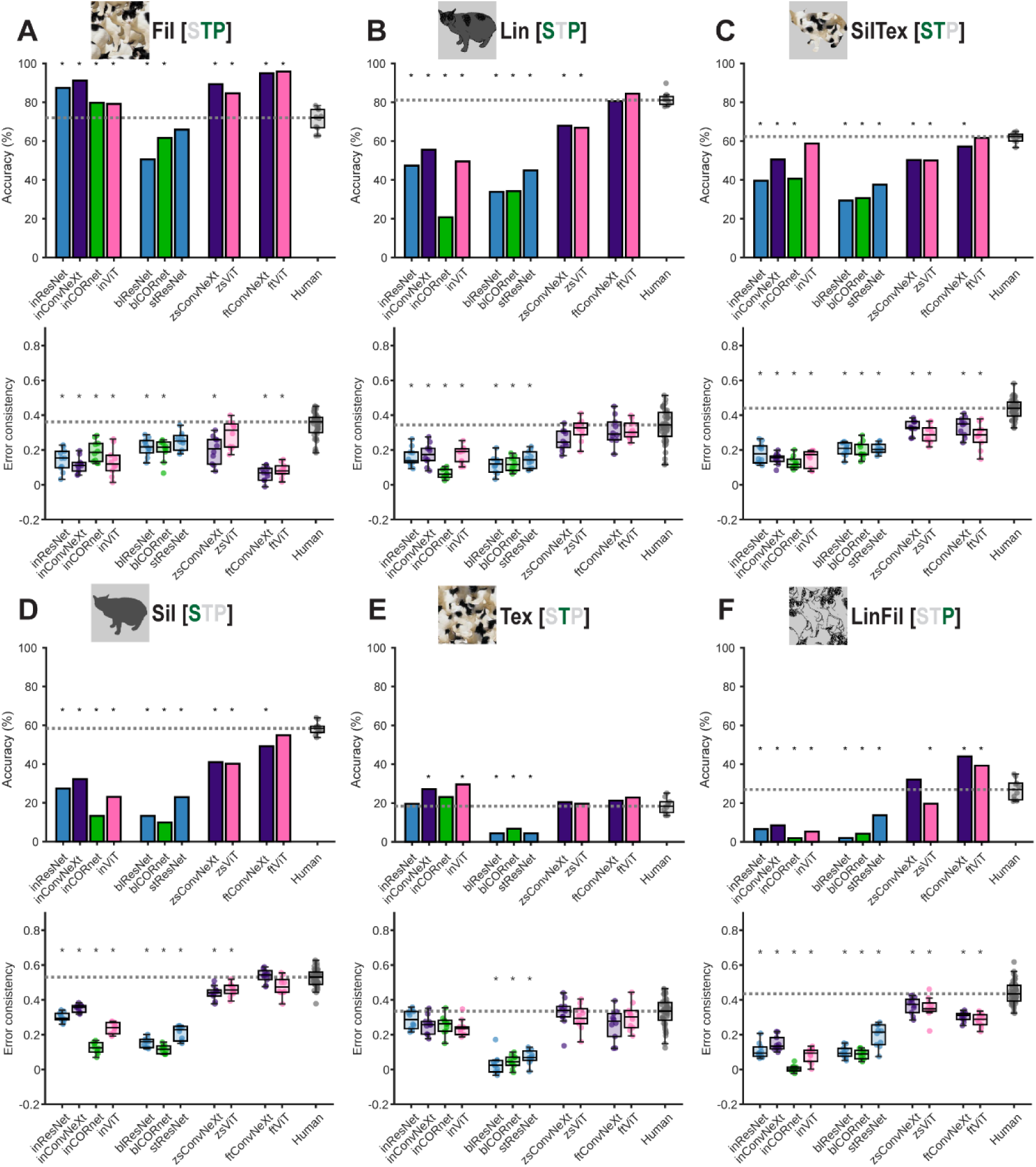
Comparison of human observers and 11 DNNs in each manipulated image condition. (**A–F**; top panels) Recognition accuracy of the 11 DNNs (listed in Fig. 1C) and humans in each condition. For details on image conditions, see Fig. 1A and text. For the human box plots, overlaid dots represent the accuracy of each pooled subject (N = 10 after pooling; see Methods for details). The horizontal dashed line indicates the human median accuracy. Stars above the bars denote significant differences between DNN accuracy and human accuracy (two-sided Wilcoxon signed-rank tests; p < 0.05, Bonferroni-corrected; exact statistics in the supplementary Excel file). (**A–F**; bottom panels) Human-to-DNN error consistency for the 11 DNNs and human-to-human error consistency in each condition. Overlaid dots for DNNs represent error consistency between that DNN and each individual pooled subject, and, for humans, between each pair of pooled subjects. The horizontal dashed line indicates the median human-to-human error consistency. Stars above DNN box plots indicate significant differences between DNN-to-human error consistency and human-human error consistency (bootstrap tests; p < 0.05; Bonferroni-corrected; see Methods for details; exact statistics in the supplementary Excel file).

### ImageNet-trained models all diverged from human cue reliance

We first compared the recognition behavior of two standard ImageNet-trained CNNs (inResNet and inConvNeXt) with humans across the six manipulation conditions. In addition to overall accuracy, we compute DNN-to-human error consistency, which measures the extent to which models and humans make mistakes on the same images, providing an image-wise assessment of human-likeness. If a model struggles with images that humans find easy or *vice versa*, it is not human-like even if the overall accuracy is similar. By comparing DNN-to-human error consistency against the distribution of error consistency among human observers, we evaluated whether DNN-to-human alignment deviated from typical inter-subject variability. Differences in recognition accuracy or error consistency between each DNN and human observers were assessed with a two-sided Wilcoxon signed-rank test and a non-parametric bootstrap test, respectively, applying Bonferroni correction across the eleven models (see Methods for details; exact statistics in supplementary Excel file).

Both CNNs significantly outperformed humans in the **Fil** condition (**Fig. 2A**; top). In the **Tex** condition, no significant difference was observed for inResNet, while inConvNeXt exhibited significantly higher accuracy than humans (**Fig. 2E**; top). In the remaining four conditions, both CNNs significantly underperformed humans (**Fig. 2B–D, F**; top). Human-model error consistency for both CNNs was significantly lower than error consistency between human subjects in all conditions except **Tex** (**Fig. 2A–F**, bottom). Collectively, the two CNNs matched human behavior only when texture was the sole available cue, confirming the earlier observation that standard CNNs rely heavily on the texture cue^20,23^.

We next investigated whether ImageNet-trained DNNs with alternative architectures better matched human behavior. A fundamental difference between CNNs and the primate ventral stream is that CNNs operate in a purely feedforward manner, whereas the ventral stream has abundant lateral and feedback connections. Given the evidence that recurrent processing contributes to object recognition^37–41^, we tested whether ImageNet-trained CORnet-S (inCORnet), a popular recurrent CNN^33^, would exhibit higher behavioral alignment with humans. However, inCORnet either fell further below human levels than the CNNs (**Lin**, **Sil**, **LinFil**; **Fig. 2B, D, F**) or showed no notable change (**Fil**, **SilTex**, **Tex**; **Fig. 2A, C, E**).

Vision Transformers (ViTs) represent another promising approach given their state-of-the-art performance on classification benchmarks^11,42^. ViTs use the same self-attention mechanisms as large language models (LLMs) to perform spatial reasoning by drawing relationships between different patches within the same image. Although ImageNet-trained ViT (inViT) more closely approximated human accuracy in the **SilTex** condition, its error consistency remained low (**Fig. 2C**). In other conditions, we observed no improvements relative to the CNNs.

Taken together, all ImageNet-trained DNNs, regardless of architecture (including feedforward CNNs, recurrent CNN, and vision transformer), continue to have a large performance gap and/or a large behavioral misalignment with humans in all conditions except when texture is the sole available information. In addition, when both texture and internal parts are available (**Fil**), all four models outperform humans but have low behavioral alignment to humans.

### Blurred and stylized training did not improve human alignment

Beyond architecture, the training diet critically shapes DNN behavior. Prior studies showed that training on blurred versions of ImageNet increased a model’s shape bias when tested on shape-texture cue-conflict images and enhanced the alignment of internal model representations with ventral stream neural activity as measured by fMRI^32^. Similarly, CNNs trained on a stylized version of ImageNet also achieved a greater shape bias as measured by shape-texture cue-conflict images^20^. However, a higher shape bias measured by these cue-conflict images could reflect either better utilization of the shape cue or a worse ability to utilize the texture cue, since the two cues are directly pitted against each other. Thus, it remains unknown how exactly these DNNs’ cue-reliance profiles align with humans. To fill these gaps, we tested blurred-image-trained ResNet50 (blResNet)^32^, blurred-image-trained CORnet-S (blCORnet)^32^, and stylized-image-trained ResNet50 (stResNet)^20^ on our image set wherein different cues were independently manipulated.

For blurred-image-trained models, in the **Fil** condition where ImageNet-trained models outperformed humans, both blResNet and blCORnet instead significantly underperformed humans, with slight improvement in error consistency (**Fig. 2A**). In the **Lin**, **SilTex**, **Sil**, and **LinFil** conditions, where ImageNet-trained CNNs significantly underperformed humans, the accuracy of blResNet and blCORnet became even worse (**Fig.2B–D, F**; top). In the **Tex** condition, although standard CNNs mostly matched humans, blResNet and blCORnet significantly underperformed humans and had low error consistency to humans (**Fig. 2E**). The stylized-image-trained model, stResNet, performed slightly better than blurred-image-trained models, but did no better than ImageNet-trained models and performed substantially below humans in 5 out of 6 conditions (all except **Fil**) with low error consistency to humans.

Therefore, training on blurred or stylized images failed to enhance human-like recognition behavior and was often counterproductive, yielding worse performance and/or human alignment compared to standard ImageNet-trained models.

### Fine-tuned CLIPs nearly closed the gap in shape-preserved conditions

CLIP is a self-supervised learning approach that associates images with natural language captions, moving beyond models specialized for a single visual task^42^. CLIP models have demonstrated remarkable zero-shot generalization to classification tasks without additional retraining, and high robustness to noise and style changes, suggesting potential human-like cue usage^11^. To assess this, we evaluated two zero-shot CLIPs (zsCLIPs), one with a CNN (ConvNeXt) backbone (zsConvNeXt) and another with a ViT backbone (zsViT), on our image set. These DNNs are zero-shot because they have never been directly trained for image classification.

Compared to standard CNNs, both zsCLIPs moved closer to the human level in terms of accuracy and error consistency in the **Lin**, **Sil**, and **LinFil** conditions (**Fig. 2B, D, F**), while maintaining human-like behavior in the **Tex** condition (**Fig. 2E**). In the **Fil** and **SilTex** conditions, error consistency also increased, whereas accuracy was similar to standard CNNs (**Fig. 2A, C**). Despite these gains, both zsCLIPs still showed significant differences from humans in accuracy or error consistency in all conditions except **Tex**, indicating they only partially closed the behavioral gap.

Although zsCLIPs perform well at object classification without explicit categorization training, prior work has shown that fine-tuning on an image-classification task further improves accuracy^43^. Accordingly, we evaluated fine-tuned versions of both ConvNeXt- and ViT-based CLIP (ftConvNeXt and ftViT). Across all conditions except **Tex**, fine-tuning boosted accuracy relative to zsCLIPs. This accuracy increase brought ftCLIPs into near-alignment with human behavior in shape-preserved conditions (**Lin**, **SilTex**, and **Sil**; **Fig. 2B–D**). In contrast, in the **Fil** condition, where standard CNNs already exceeded human accuracy, ftCLIPs’ performance rose even further above the human level and error consistency dropped to the lowest among all models tested thus far (**Fig. 2A**). In the **LinFil** condition, ftCLIPs significantly outperformed humans with below-human error consistency whereas most other DNNs underperformed (**Fig. 2F**). Together, while ftCLIPs relatively well replicated human recognition in shape-preserved conditions, they amplified over-robustness when the global shape information was disrupted either alone (**Fil**) or together with texture (**LinFil**).

In summary, regardless of architecture (CNN or ViT), Contrastive Language-Image Pre-trained (CLIP) models that are fine-tuned on classification tasks had the highest overall alignment to humans. However, when the global shape cue was disrupted, fine-tuned CLIPs often outperformed humans with a low error-pattern alignment to humans. In these conditions, zero-shot CLIPs are slightly better aligned to humans.

### A systematic evaluation of >200 DNNs reveals systematic human-DNN differences

To confirm the generalizability of these findings, we expanded our analysis to include 198 additional DNNs from the Brain-Score database^44,45^, covering a wider range of architectures, training diets, and training approaches (Table S1). In shape-preserved conditions (**Lin**, **SilTex**, **Sil**), the models whose behavior most closely matched humans were predominantly ftCLIP variants (**Fig. 3B–D**; red-outlined dots and pink/purple squares; Table S2). By contrast, when shape information was absent (**Fil**, **Tex**, **LinFil**), ftCLIPs tended to surpass human accuracy but had low error consistency to humans (**Fig. 3A, E, F**; red-outlined dots and pink/purple squares). Consequently, the models most similar to humans in these conditions were not dominated by ftCLIPs (Table S2). In these three shape-absent conditions, although the model closest to human behavior differed across conditions, zsCLIPs consistently remained relatively close to human behavior, with zsViT ranking among the top 10 in all three conditions (**Fig. 3A, E, F**; pink diamonds; Table S2). Thus, ImageNet fine-tuning is unnecessary or even counterproductive for achieving human-like behavior when shape cues are unavailable.

**Fig. 3.**
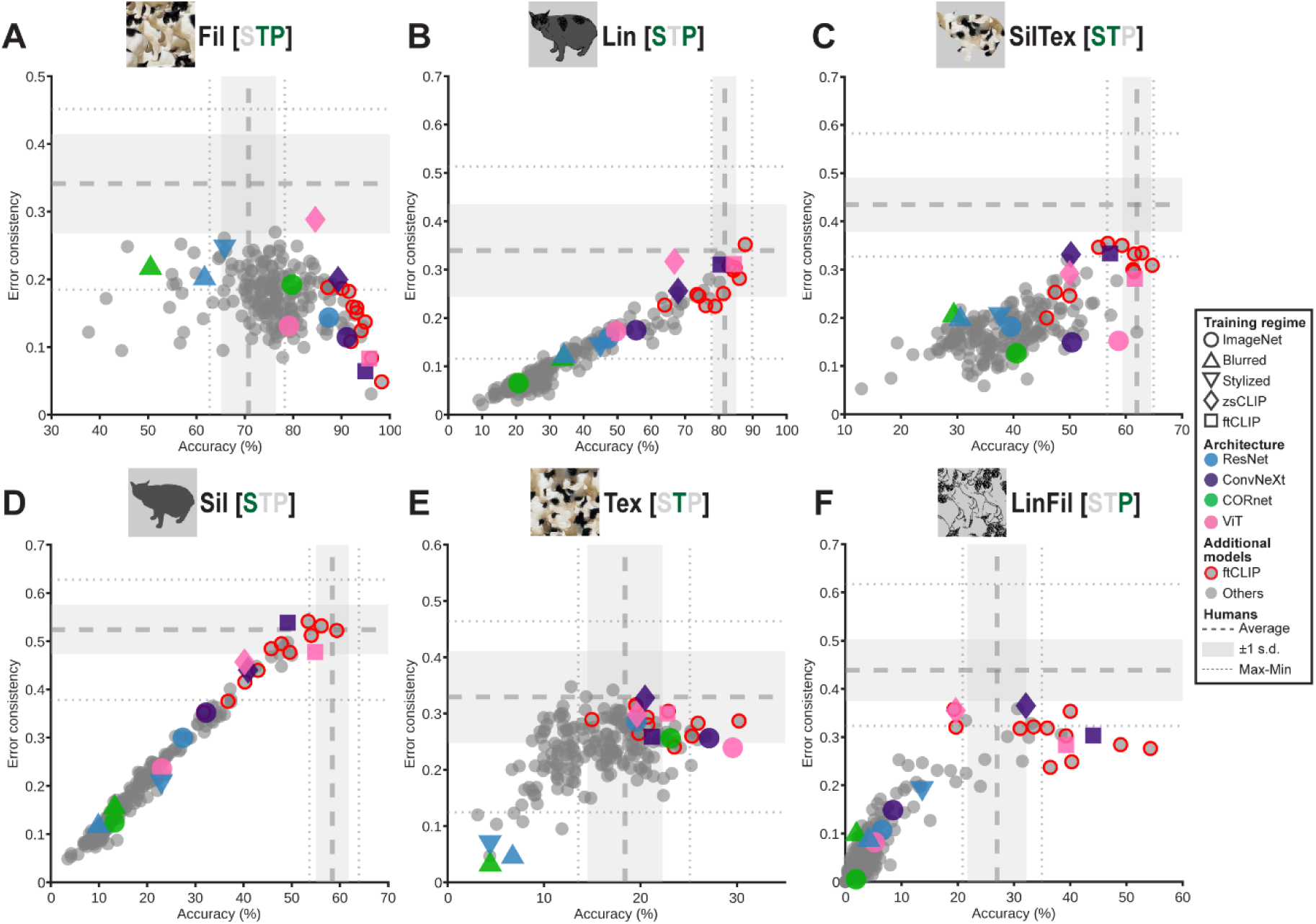
Large-scale evaluation of 209 DNNs. (**A–F**) Scatter plots of 209 DNNs’ accuracy (x-axis) against average error consistency with humans (y-axis) under each image condition. Horizontal and vertical dashed (/dotted) lines denote the average (/min-max) human accuracy and human-to-human error consistency, respectively. The gray shaded regions denote the mean ± 1 s.d. of human accuracy and human-human error-consistency. The 11 DNNs from earlier analyses are highlighted with distinct markers and colors as per the inset; other ftCLIPs are shown as gray dots outlined in red; remaining models are gray dots without outlines. The ten models whose behavior most closely matches humans in each condition are listed in Table S2.

In addition to all DNNs in the Brain-Score database, we also tested an additional model, Stable Diffusion classifier, a generative model trained with noise-added images^29^. This model was previously found to exhibit a human-like shape bias when tested on texture-shape cue-conflict images, and had high accuracy and error consistency with humans on out-of-distribution images^29^. However, Stable Diffusion classifier did not behave human-like on our stimuli set (**Fig. S3**): its performance was exceptionally low in texture-preserved conditions (**Fil, SilTex, Tex**) and only mid-rank among the tested DNNs in the other conditions (**Lin, Sil, LinFil**). This suggests that this model has a compromised ability to utilize the texture cue, and its high shape bias as assessed by shape-texture cue-conflict images is likely artifactual—due to an inability to use the texture cue instead of a genuinely improved ability to utilize the shape cue.

Thus, a systematic evaluation of >200 DNNs in the Brain-Score database revealed that no single model replicated human recognition behavior both when the global shape information is present and when it is missing: ftCLIP models excel when the global shape cue is available, whereas other models— such as zsCLIPs—better approximate human behavior when global shape is absent.

A striking observation in this analysis is the divergent relationship between accuracy (quantifying overall performance) and error consistency (quantifying error alignment with humans) in shape-preserved versus shape-absent conditions. In shape-preserved conditions (**Lin**, **SilTex**, **Sil**), improvements in model accuracy coincided with increases in error consistency, and human accuracy remains the approximate upper bound on accuracy (**Fig. 3B–D**). In contrast, under shape-absent conditions, multiple models surpassed human accuracy, and higher accuracy did not reliably translate into higher error consistency with humans (**Fig. 3A, E, F**). This pattern suggests that humans disregard certain non-shape cues that models nonetheless leverage for classification tasks, allowing those models to classify images that are challenging for humans.

In addition, our earlier analysis of eleven DNNs found that models previously suggested to better predict neural activity in the ventral visual stream (CORnet-S and blurred-image-trained models) did not exhibit more human-like behavior (**Fig. 2**). To examine this on a larger scale, we correlated each DNN’s behavioral similarity to humans with its ability to predict neural activity in the inferotemporal (IT) cortex, as measured in Brain-Score^44,45^. We used three behavioral similarity metrics: the absolute difference in the accuracy between humans and each model (|Δaccuracy|), the difference between human-to-human and human-to-DNN error consistency (Δerror-consistency), and the root-sum-square of these two metrics after normalization (for details, see Methods). A negative correlation is expected if IT predictability is associated with greater human alignment (i.e., smaller human-DNN behavioral distance corresponds to higher IT predictability). However, across all six conditions and three behavioral similarity metrics, we found no significant correlations, indicating that higher IT predictability as measured by the Brain-Score dataset does not predict more human-like recognition behavior (**Fig. S4**, top-left magenta annotations). Note that the image set used for IT recordings in the Brain-Score database differs substantially from the image set used to compute behavioral similarity in our study, which warrants cautious interpretation (see Discussion).

Brain-Score also includes numerous behavioral benchmarks, each of which measures DNN-human behavioral alignment using different metrics across a wide range of image sets. Thus, we further assessed how human alignment in each condition of our image set relates to the overall human alignment measured by existing Brain-Score behavioral benchmarks. Specifically, for each image condition, we computed the correlation between the average score across all Brain-Score behavioral benchmarks and the three behavioral similarity metrics used above. Again, a *negative* correlation is expected if the Brain-Score benchmark predicts human-likeness in our image set. We found that the **Lin**, **SilTex**, **Sil**, and **LinFil** conditions showed relatively strong negative correlations across all three metrics (**Fig. S4**, top-right blue annotations; r = −0.48 to −0.62, all ps < 0.05), suggesting that these conditions have overlaps with existing benchmarks. By contrast, the **Tex** condition showed weaker correlations (r = −0.09 to −0.13; n.s. for |Δaccuracy|), suggesting that the Brain-Score benchmarks are less informative for object recognition when texture information alone is available. Interestingly, the **Fil** condition showed a *positive* correlation between our three *distance* metrics and Brain-Score behavioral benchmarks, suggesting that existing benchmarks do not capture our **Fil** condition at all. Together, these results suggest that our image set has both overlap and differences compared to the existing behavioral benchmarks in Brain-Score.

### All DNNs underperformed humans when the global shape was decoupled from the local shape

When only the global shape cue was available (**Sil**), some ftCLIPs perfectly replicated humans’ recognition behavior (**Fig. 3D**). However, previous work showed that the silhouette recognition performance of a standard ImageNet-trained CNN catastrophically drops when jagged edges were added to the silhouette contour, even though humans remain nearly unaffected^21^. This raises the question of whether ftCLIPs recognize silhouettes in the same way as humans do and are robust to local distortions of the global shape.

To test this and evaluate various DNN models on global shape-based perception that is fully decoupled from local cues, we conducted Experiment 2. We rendered silhouettes as assemblies of small cross marks at a low (**Sparse**) or high (**Dense**) density to disrupt local shape while preserving the global shape (**Fig. 4A**). Human recognition for the **Sil**, **Sparse**, and **Dense** conditions was collected via another online experiment with 10 new subjects, using the same task design as Experiment 1 (**Fig. 1B**; for details, see Methods). To evaluate the effect of decoupling local and global shape cues regardless of baseline performance on silhouette images, for each DNN and human subject we excluded any images misclassified in the **Sil** condition from subsequent analyses.

**Fig. 4.**
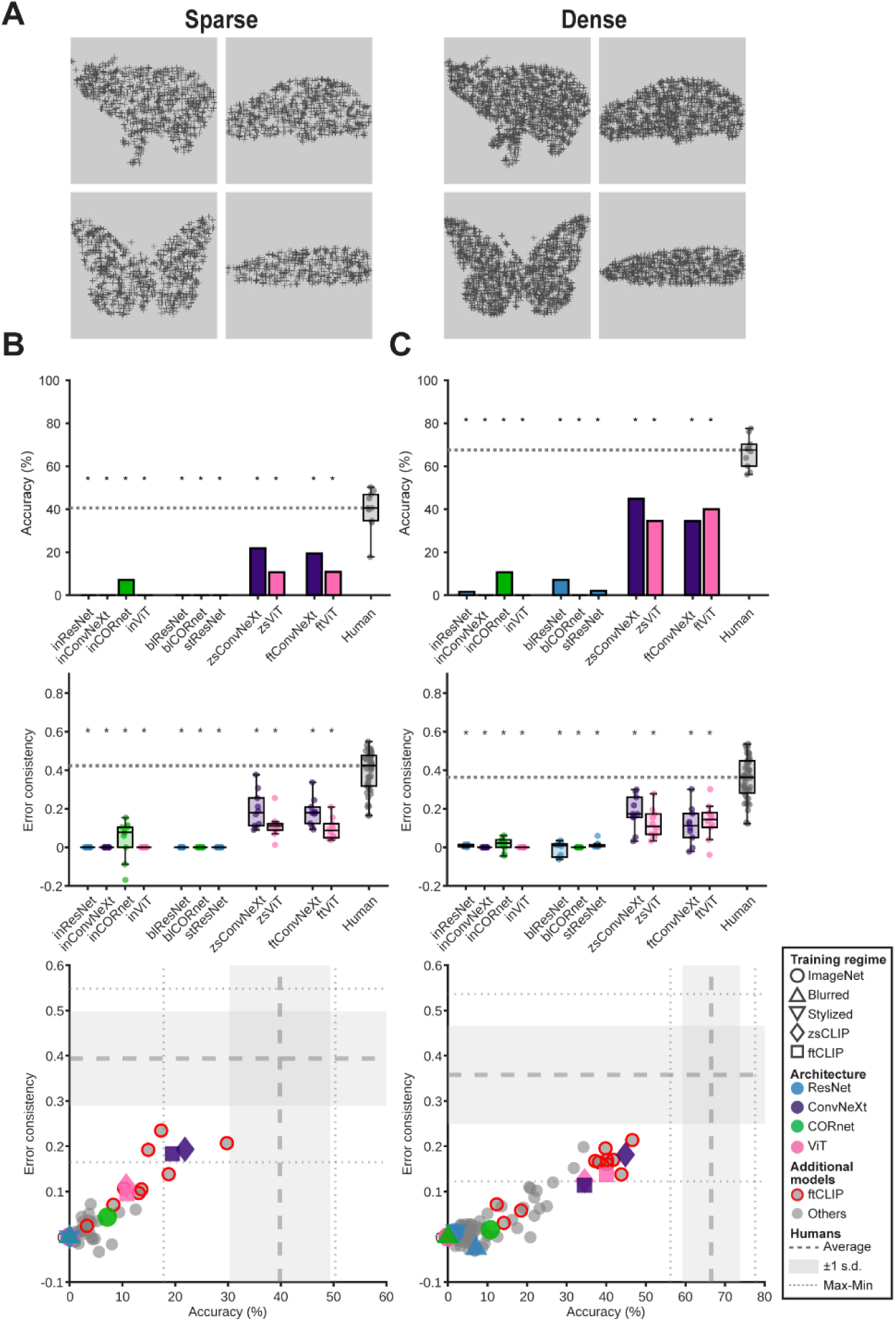
Comparison of human and DNN recognition behavior when local and global shape cues are decoupled (Experiment 2). **(A)** Example images for the Sparse and Dense conditions. Corresponding original images are shown in Fig. 1A. (**B–C**; top panels) Recognition accuracy of the 11 DNNs and human subjects in Sparse **(B)** and Dense (C) conditions. For human box plots, the overlaid dots represent the accuracy of each subject (N = 10). The horizontal dashed line indicates the median human accuracy. Stars above the bars denote significant differences between a DNN’s accuracy and human accuracy (two-sided Wilcoxon signed-rank tests; p < 0.05; Bonferroni-corrected; exact statistics in the supplementary Excel file). (**B–C**; middle panels) Human-to-DNN error consistency for the 11 DNNs and human-to-human error consistency. The horizontal dashed line indicates the median human-to-human error consistency. Stars above DNN box plots indicate significant differences as compared to humans (bootstrap tests; p < 0.05, Bonferroni corrected; exact statistics in the supplementary Excel file). (**B–C**; bottom panels) Scatter plots of 209 DNNs’ accuracy (x-axis) against average error consistency with humans (y-axis). The format is the same as Fig. 3.

We first evaluated the same eleven models from **Fig. 2**. No models reached human-level accuracy or error consistency in either the **Sparse** or **Dense** condition (**Fig. 4B, C**; top and middle). zsCLIPs and ftCLIPs outperformed other models, suggesting that CLIP yields the most robust recognition performance based on the global shape information. inCORnet (i.e., ImageNet-trained CORnet) is the only other DNN that exhibited above-chance accuracy in both conditions, but with accuracy and error consistency far below the human level. Other models had 0% or near 0% accuracy, indicating that they were unable to recognize an object based on the global shape when it is decoupled from local shape.

To better understand the drastic failure of DNNs, we examined the predicted classes for **Sparse** and **Dense** images. Most predictions were confined to a small number of classes featuring cross-like patterns, such as “crossword puzzle,” “jigsaw puzzle,” or “window screen” (Table 1). For example, inConvNeXt labeled 95% of **Sparse** images as “crossword puzzle,” and even CLIP models often predicted “chain mail”, “window screen” or “nematode”. This confirms that DNNs commonly ignore the global shape and are strongly biased toward local shape cues.

**Table 1.** Top 1 predicted classes assigned to more than 10% of Sparse and Dense exemplars; percentages in parentheses indicate the proportion of exemplars predicted as that class.

Sparse
| **inResNet** | **inConvNeXt** | **inCORnet** | **inViT** |
| --- | --- | --- | --- |
| crossword puzzle (72.5) | crossword puzzle (95.3) | jigsaw puzzle (38.6) | window screen (83.0) |
| strainer (11.7) | – | screw (19.9) | – |
| – | – | envelope (11.7) | – |
| **blResNet** | **blCORnet** | **stResNet** | **–** |
| stole (24.6) | hair slide (33.9) | envelope (42.1) | – |
| chain mail (11.7) | jigsaw puzzle (18.7) | strainer (16.4) | – |
|  | dishrag (15.8) | theater curtain (11.1) | – |
| **zsConvNeXt** | **zsViT** | **ftConvNeXt** | **ftViT** |
| chain mail (39.2) | chain mail (19.9) | nematode (24.0) | nematode (28.1) |
| plane (12.3) | maze (18.1) | window screen (10.5) | crossword puzzle (19.3) |
| – | knot (14.0) | – | – |

Dense
| **inResNet** | **inConvNeXt** | **inCORnet** | **inViT** |
| --- | --- | --- | --- |
| window screen (46.8) | crossword puzzle (54.4) | chain mail (37.4) | nematode (55.0) |
| jigsaw puzzle (21.1) | coral fungus (20.5) | stole (21.1) | window screen (36.8) |
| **blResNet** | **blCORnet** | **stResNet** | **–** |
| envelope (33.3) | stole (19.3) | hair slide (24.0) | – |
| strainer (14.6) | chain mail (18.7) | chain mail (18.1) | – |
| **zsConvNeXt** | **zsViT** | **ftConvNeXt** | **ftViT** |
| plane (11.7) | – | nematode (14.0) | – |

We then evaluated the additional 198 DNNs as used in **Fig. 3** (Table S1). Although CLIP models generally outperformed other DNNs regardless of their architecture, the best model’s accuracy and error consistency remained far below the average human level in both conditions (**Fig. 4B, C**; bottom; Table S2). Stable Diffusion classifier had near-0 (1.41%) accuracy in the **Sparse** condition and low (12.7%) accuracy in the **Dense** condition (**Fig. S3**). In addition, similar to results in Experiment 1, we did not observe a significant correlation between neural predictability score based on ventral stream neural recordings (as measured in Brain-Score) and closeness to human behavior (**Fig. S4**, rightmost column, magenta annotations). By contrast, mean Brain-Score behavioral benchmarks exhibited significant correlations with human-alignment in our dataset (**Fig. S4**, rightmost column, blue annotations).

To examine the possibility that the DNNs might have produced multiple answers based, respectively, on the local and global shape information, we repeated the above analysis using DNNs’ top-5 predictions instead of the top-1 prediction. In this analysis, a response was counted as correct if the correct label appeared among the top five predictions. Although some CLIP models that already showed relatively high accuracy in the original analysis improved to near-human levels, many other models remained at very low accuracies (**Fig. S5A**). This analysis further confirms the catastrophic failure of non-CLIP models to recognize objects based on the global shape in the **Sparse** and **Dense** conditions.

Intuitively, the difficulty of recognizing **Sparse**/**Dense** images depends on the sizes at which the images are presented: the smaller the images, the easier it should be to recognize the object based on the global shape. In our human experiments, all images were presented centrally at 10° × 10° of visual angle, following standard “core object recognition” tasks^1,46^. However, DNNs do not have a foveal bias and the object size presented to the DNN is arbitrary (we used images from the ImageNet database, at 256×256 pixels). To confirm that image size did not substantially affect our results, we assessed DNN performance on modified **Sparse** and **Dense** images, in which the objects were half the sizes compared to the original images (**Fig. S5B**, left). This manipulation renders the global shape more salient. Yet, when presented with these half-sized images, there were no substantial improvements in model performance (**Fig. S5B**, right). This result suggests that our results are robust across image sizes.

Together, these findings suggest that, compared to humans, all DNNs consistently exhibit a bias toward utilizing the local shape cues over global shape cues, regardless of architecture, training diet, or training objective, although CLIP models generally did the best. Crucially, Experiment 2 also reveals a large and persistent behavioral gap between all DNNs and humans when the global shape is the only information available for object recognition and is decoupled from local shape cues.

### Activations evoked by local shape cues masked those evoked by the global shape in DNNs

To shed light on the catastrophic performance of all DNNs relative to humans in Experiment 2, where global shape was decoupled from local shape, we probed the models’ internal representations. There are two potential scenarios. First, DNNs might retain both global and local shape information but with local shape representation being dominant, causing outputs to reflect cross-like objects. If this is the case, multivariate decoding analysis applied to model internal representations should be able to recover the object category (defined here as basic object category, such as ‘bear’) as defined by the global shape even when the model output fails. Alternatively, the DNNs might completely lose the object information defined by the global shape; in this case, decoding accuracy based on model internal representations would be as poor as the model’s output.

To test these possibilities, we performed object category decoding in the **Sparse** and **Dense** conditions using internal representations from four representative layers of each of the 11 DNNs tested earlier (see **Fig. 1C**), covering early to late processing (**Fig. 5A**, left; Table S3). We decoded object category using a leave-one-exemplar-out approach (**Fig. 5A**, right). This analysis probes whether different exemplar images of the same object category are represented similarly. Strikingly, all models, even those with 0% recognition accuracy, showed significant above-chance decoding accuracy from the middle activation layer onward in both conditions (ps < 0.05; label permutation test; Bonferroni corrected across layers; **Figs. 5B** and **S5C–E**). Category decoding accuracy increased in deeper layers. This suggests that the poor performance of the DNNs arises not from a strict inability to encode the global shape but from an inability to use such information in the classification output.

**Fig. 5.**
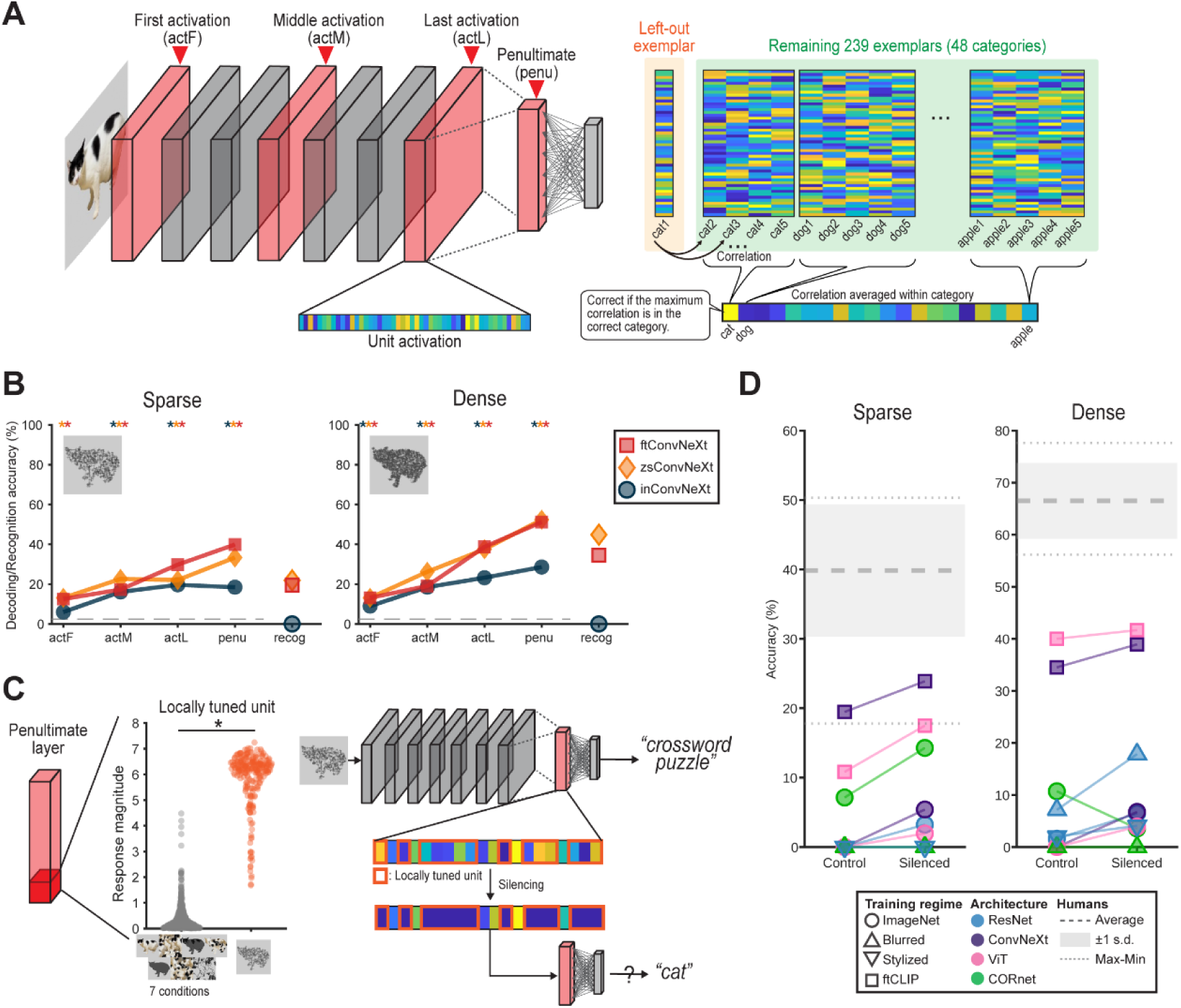
Impact of silencing locally tuned units in the Sparse and Dense conditions. **(A)** Leave-one-exemplar-out decoding schematic. For each of four representative layers (the first activation layer [actF], the middle activation layer [actM], the final activation layer [actF], and the penultimate layer [penu]), we correlated the activation pattern of the left-out exemplar with those of all other exemplars in the same condition, averaged these correlations within each category, and classified the exemplar by selecting the category with the highest mean correlation. Repeating this procedure for each exemplar yielded decoding accuracy per layer. **(B)** Decoding accuracy for the three ConvNeXt-based models at the four representative layers (actF, actM, actL, penu) under the Sparse and Dense conditions. Dashed line indicates chance (1/41; for details, see Methods). Stars indicate significant difference from chance (label permutation test; p < 0.05; Bonferroni-corrected over layers; see Methods for details; exact statistics in the supplementary Excel file). The rightmost “recog” column shows recognition accuracy reproduced from Fig. 4 for comparison. Equivalent results for other architectures are in Fig. S5C–E. **(C)** Left: Response magnitude of a representative inResNet unit in the penultimate layer whose activation differed significantly between Sparse/Dense and the 7 conditions used in Experiment 1 (two-sample t-test, p < 0.05; Bonferroni-corrected across all units in the same layer). Right: Illustration of the silencing procedure, in which all locally tuned units’ activations were set to zero in the penultimate layer, before passing to the final fully connected layer. **(D)** Recognition accuracy of the 11 DNNs in the Sparse (left) and Dense (right) conditions before and after silencing the locally tuned units. DNNs are coded by markers and colors per the inset. The horizontal dashed line and dotted lines denote the mean and range of human accuracy; the gray shaded region denotes the mean ± 1 s.d. of human accuracy.

We reasoned that if this is indeed the case, silencing units that are responsive to the local shape cues should unmask the global shape information and improve performance. To test this, for each unit in the penultimate layer, we assessed whether its response to **Sparse** or **Dense** images differed significantly from its response to all 7 conditions in Experiment 1. Since a significant difference suggests a non-specific response to local cross shapes regardless of the global shape, we defined such units as locally tuned units (**Fig. 5C**; left). We then examined the DNNs’ predictions after silencing these units by setting their activations to zero before passing through the final fully connected layer (**Fig. 5C**; right). We excluded zsCLIPs from this analysis because their predictions are based on the similarity between a test image and text embeddings (for details, see Methods).

We found that a substantial proportion of penultimate-layer units were tuned to local cross structures in both **Sparse** and **Dense** conditions (34.77–79.30% for **Sparse**, 28.87–78.19% in **Dense**; **Fig. S5F**). Remarkably, after silencing these large portions of units, many models showed improved accuracy. In the **Sparse** condition, six out of nine models had increased accuracy, while the remaining three (blResNet, blCORnet and stResNet) remained at zero (**Fig. 5D**; left). In the **Dense** condition, seven models improved, whereas inCORnet showed decreased accuracy and blCORnet remained unchanged (**Fig. 5D**; right). These improvements suggest that the dramatic DNN failure in the **Sparse** and **Dense** conditions arise partly because global shape representations are overshadowed by the dominant local-cross activations in the penultimate layer. However, even after this intervention, all DNNs’ accuracy remained far below the average human level, suggesting that local-signal masking is not the sole cause of their poor performance and deficiencies in extracting the global shape information likely also contribute to this persistent AI-human gap.

### Better models contained richer object information in their later layers

Our results thus far reveal that training exerts a greater influence on DNN recognition behavior than architecture: in both experiments, identically structured models exhibited markedly different robustness to image perturbations depending on their training. To understand how these training-dependent differences emerge, we investigated internal representations of the 11 DNNs (**Fig. 1C**) in response to images employed in Experiment 1 by probing four representative layers from each DNN covering early to late processing (**Fig. 5A**, left).

We first used representational similarity analysis (RSA) to quantify shape, texture and internal part-related information in each layer. To this end, we created model RDMs (**Fig. S6A**) for shape, texture, or internal-parts information based on the 7 conditions (**Ori** and 6 manipulation conditions), which were correlated with empirical RDMs estimated from the activation patterns of each DNN layer. This correlation was performed for each exemplar image, with the result averaged across 240 exemplar images.

Across all 11 DNNs, visual feature representations exhibited dynamic changes across the network hierarchy, with broadly similar trajectories (**Fig. S6B–E**). We therefore focused on differences in the penultimate-layer representation between DNNs with the same architecture but different training. In ConvNeXt-based models (**Fig. S6B**), texture representation in CLIP models (zsConvNeXt and ftConvNeXt) significantly reduced relative to the ImageNet-trained model (inConvNext vs. zsConvNeXt: t_239_ = 10.72, p = 1.41×10^−21^; inConvNext vs ftConvNext: t_239_ = 16.95, p = 2.97×10^−42^; paired t-test; Bonferroni corrected), and internal-parts representation in ftConvNeXt increased compared to the other two models (ftConvNext vs. inConvNeXt: t_239_ = 15.07, p = 6.26×10^−36^; ftConvNext vs. zsConvNext: t_239_ = 21.87, p = 2.24×10^−58^). ViT-based models (**Fig. S6C**) exhibited similar trends, albeit of much smaller magnitudes for both texture (inViT vs. zsViT: t_239_ = 8.25, p = 4.08×10^−14^; inViT vs. ftViT: t_239_ = 2.06, p = 0.16) and internal parts (ftViT vs. inViT: t_239_ = 6.64, p = 8.28×10^−10^; ftViT vs. zsViT: t_239_ = 10.85, p = 5.66×10^−22^).

Training with blurred images had comparable effects in ResNet- and CORnet-based models (**Fig. S6D, E**): texture representation decreased (inResNet vs. blResNet: t_239_ = 7.05, p = 7.69×10^−11^; inCORnet vs. blCORnet: t_239_ = 13.33, p = 4.58×10^−30^), and internal-parts representation increased relative to their ImageNet-trained counterparts (inResNet vs blResNet: t_239_ = −8.52, p = 7.09×10^−15^; inCORnet vs blCORnet: t_239_ = −16.37, p = 2.57×10^−40^). Stylized-image training yielded a similar and stronger pattern for both texture (inResNet vs. stResNet: t_239_ = 14.32, p = 1.99×10^−33^) and internal parts (inResNet vs. stResNet: t_239_ = −14.61, p = 2.17×10^−34^).

Although these results confirm that training modulates which visual features are emphasized, feature-representation strength did not reliably predict recognition accuracy when that feature was the sole available cue. For example, although recognition accuracy in the **Sil** condition, where shape is the only available cue, was higher for ftConvNeXt than inConvNeXt, shape representation was slightly but significantly lower in ftConvNeXt than inConvNeXt (t_239_ = 6.35, p = 4.27×10^−9^; **Fig. S6B**; left). Similar mismatches between feature-representation strength and cue-specific recognition accuracy were also observed in other architectures and conditions. This implies that encoding a specific visual feature faithfully does not guarantee high recognition performance based on that feature. Instead, success might depend on a DNN’s ability to extract object-relevant information and group exemplars of the same object category even when their visual features vary. For example, silhouettes of a snake can differ markedly—one might be coiled, another elongated—yet to achieve high silhouette accuracy, a model must map these different shapes onto the single concept “snake”.

To directly test this possibility, for each DNN we performed leave-one-exemplar-out object category decoding at each of the four representative layers for every image condition (using the same approach illustrated in **Fig. 5A**). In ConvNeXt-based models, decoding accuracy increased steadily from the first activation layer to the penultimate layer in all image conditions (**Fig. 6**). Moreover, models with higher recognition accuracy typically also showed higher decoding accuracy in the penultimate layer. Models with other architectures (ViT, ResNet, CORnet) exhibited a similar general correspondence between penultimate-layer decoding accuracy and recognition performance (**Fig. S7**). Together, these results suggest that models that better map limited visual cues onto correct categories exhibit more robust recognition under image disruptions; moreover, this ability is not directly linked to how faithfully those visual features are represented in the model.

**Fig. 6.**
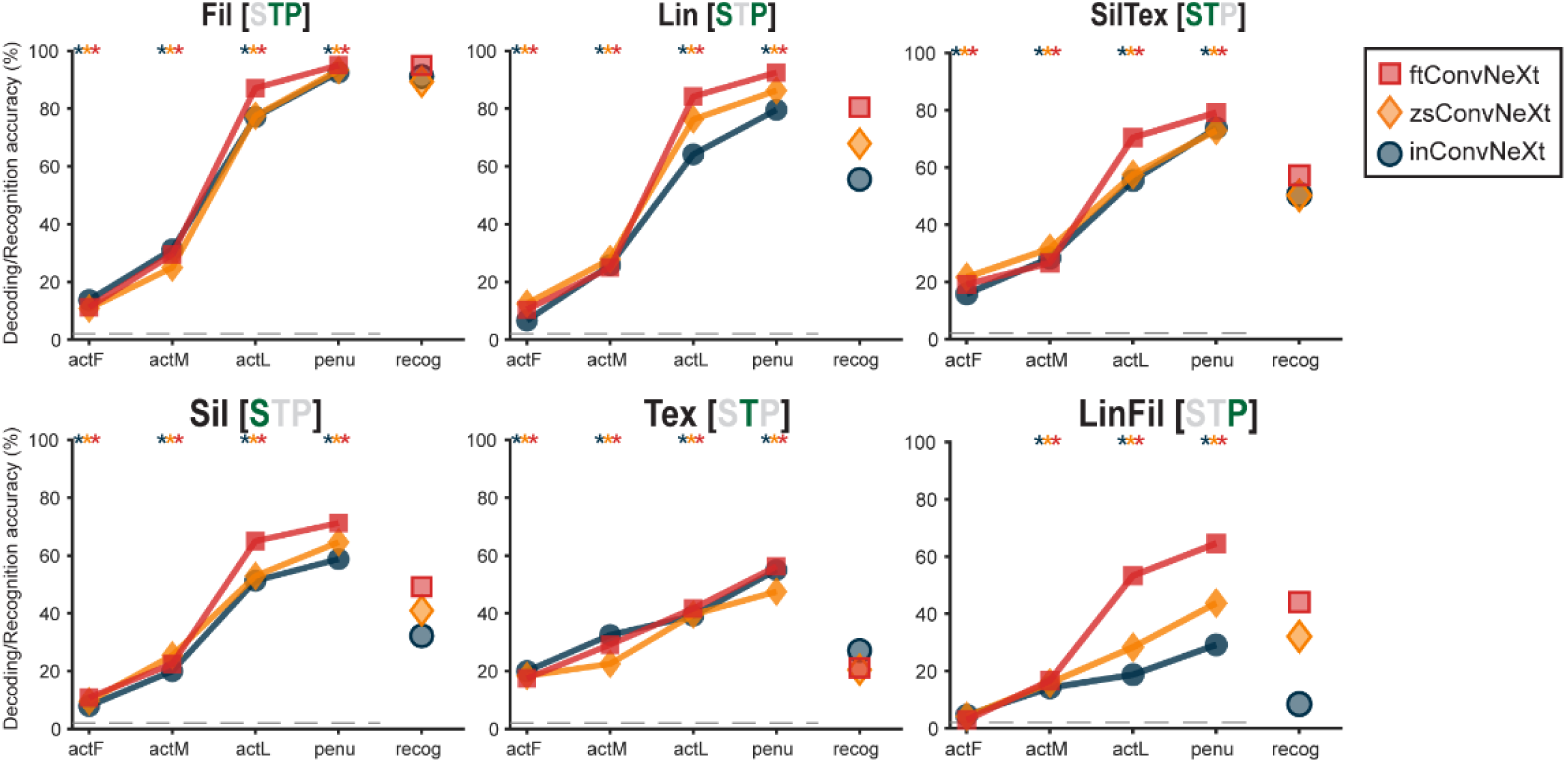
Object-category decoding from hidden-layer activations in ConvNeXt-based models. Decoding accuracy of the three ConvNeXt-based models at each of the four representative layers across conditions. Different colors denote different training approaches (red: fine-tuned CLIP; yellow: zero-shot CLIP; blue: ImageNet trained). Dashed line indicates chance level (1/48). Stars indicate significant differences from chance (label permutation test; p < 0.05; Bonferroni-corrected over layers; see Methods for details). The rightmost “recog” column shows recognition accuracy reproduced from Fig. 2 for comparison. Equivalent results for other architectures are in Fig. S7.

## DISCUSSION

A computational model that well predicts biological vision offers a powerful tool to deepen our understanding of the computational principles underlying vision. Yet, in object recognition, we have lacked a rigorous framework for evaluating the similarity of the underlying strategy leveraged by humans and each of the myriad models. Key to this question is to what extent different visual cues are relied on for object recognition, which could shed light on the sources of agreement and discrepancy between biological and machine vision. In this study, we addressed this gap by creating an image set in which different visual cues are systematically disrupted by controlled perturbations, and by comparing human observers to a wide range of DNNs. Our stimuli set differs from previous studies in perturbing each visual cue independently, which allowed us to reveal a persistent and systematic human-AI gap.

First, consistent with previous studies^20,21^, we found that standard ImageNet-trained feedforward CNNs diverged markedly from human observers when recognition depended on limited cues. However, models previously reported to improve human-alignment—recurrent CNNs^33^ and networks trained on blurred or stylized images^20,32^—showed no consistent gains. Although fine-tuned CLIP models achieved human-like performance when the global shape was preserved, they exhibited low behavioral alignment to humans when the global shape was disrupted. In these latter conditions, other models such as zero-shot CLIP emerged as the most human-like performers. Consequently, no single DNN replicated the human cue-reliance profile. Moreover, decoupling local and global shape cues revealed a persistent local-shape bias in all DNNs tested: none achieved human-level recognition based on the global shape. Finally, our RSA and decoding analyses demonstrated how training impacts hidden-layer representations and how these representational changes underpin each model family’s recognition behavior. Taken together, our image set uncovers both failures pervasive across DNNs and distinct limitations tied to particular model families.

In Experiment 1, we found that standard ImageNet-trained feedforward CNNs were highly susceptible to visual cue disruptions: they matched or exceeded human accuracy only in the **Fil** and **Tex** conditions, where texture information is present either alone or in conjunction with internal parts information; standard CNNs performed poorly in all other conditions. Overall, this pattern of findings is broadly consistent with the well-documented fragility of standard CNNs to the disruption of local or texture information^11–16^.

Recent efforts to close the gap between standard CNNs and humans on manipulated-image benchmarks (e.g., ImageNet-C^47^, shape–texture conflict^20^, and out-of-distribution dataset^13^) have leveraged specialized training and produced models that outperform standard CNNs on specific tests. However, these models—including networks trained with blurred or stylized images^20,32^ and Stable Diffusion classifiers^29^—performed no more human-like than standard CNNs on our image set. This discrepancy suggests that existing benchmarks capture only narrow facets of recognition, and a model optimized for one facet may underperform on others. In particular, we found that models with a high shape bias as assessed by shape-texture cue-conflict images^20^ often suffered dramatic accuracy drops in our **Tex** condition and performed far below the human level (i.e., blResNet, blCORnet, stResNet, and Stable-diffusion). This suggests that their increased shape bias does not reflect a genuine prioritization of shape but rather stems from a reduced ability to utilize texture information. Indeed, by pitting shape and texture cues against each other, a high “shape bias” as evaluated by shape-texture cue-conflict images incentivizes a worse ability to utilize the texture cue and a better ability to utilize the shape cue similarly. Analysis of the models’ internal representations further supports this interpretation, showing that these models primarily discard texture information instead of enhancing shape encoding. Together, these findings expose a caveat of a commonly used benchmark and underscore the importance of systematically isolating and assessing the diagnostic value of individual visual cues.

In addition, we found that models previously suggested to exhibit higher biological plausibility or ventral stream neural alignment, including CORnet-S and blurred-image-trained CNNs^32,33,37^, did not exhibit more human-like cue usage. Likewise, our large-scale analysis of >200 DNNs revealed a lack of correlation between IT predictability and behavioral similarity to humans, echoing earlier reports of null associations between neural predictability and object recognition performance^1,18,48^. We propose two explanations here. First, the observed results may be attributable to the mismatch between the image sets used to assess behavioral alignment (our image set) and neural alignment (the existing image sets in Brain-Score). The Brain-Score database, widely used to assess DNNs’ alignment to neural recordings, currently primarily includes neural responses to “core object recognition” tasks, where clear object images are used. However, the primate brain is endowed with evolutionally advantageous mechanisms that facilitate object recognition under challenging conditions, such as low light, cluttered, or ambiguous viewing conditions (e.g., to navigate at dusk and avoid hidden predators). Thus, broader metrics for assessing neural alignment using a variety of tasks and images may allow the neural predictability of a DNN to better predict its behavioral alignment with humans. Second, most neural recordings in the Brain-Score database are from the ventral visual stream, and here we only investigated neural predictability based on recordings from IT cortex—the final stage of ventral stream processing. Yet, growing evidence points to significant contributions from brain regions outside the ventral stream to object recognition: recent fMRI studies suggest that ventral stream responses alone may not be able to account for shape perception^23,49^; the prefrontal cortex is implicated in challenging recognition tasks^41,50–52^; and studies have also revealed object information in the dorsal visual stream and prefrontal regions^53–56^. To what extent these extra-ventral-stream regions contribute to object recognition and their respective computational roles remain unclear. Nonetheless, the present findings suggest that in order to achieve a truly human-aligned model for object recognition, our modeling targets likely will need to be expanded to include these additional neural substrates.

Our results highlight the unique strengths of CLIP models regardless of architecture, with the remaining gaps indicating directions for future improvement. First, zsCLIP models were generally more human-like than ImageNet-trained models, which may indicate that natural-language supervision—loosely paralleling how human children learn about the visual world—is important for acquiring robustness closer to human-level performance. Importantly, models sharing the same architecture exhibited markedly different recognition patterns depending on whether they were trained for image categorization or with CLIP’s natural language supervision, suggesting that how a model is trained has a greater influence on its recognition strategy than its architecture. These observations resonate with recent reports of human-like high-level object concepts emerging in models trained with natural language supervision, regardless of backbone architecture^30,31,57^.

Second, fine-tuning on ImageNet made CLIP models perfectly match human behavior when the global shape was preserved, suggesting that additional training on a categorization task can be beneficial. However, this gain came at the cost of reduced human-likeness in shape-absent conditions, especially for the **Fil** condition, where ftCLIPs often exceeded human accuracy and showed markedly lower error consistency with humans. We suspect that this drawback arises because ImageNet’s fine-grained categories—over 100 dog breeds and more than 50 bird species—encourage discrimination based on subtle patterns or parts that humans typically do not rely on (such as feather patterns in birds). In other words, the divergence from human behavior could be attributable to training on a task that humans are unable to solve in the same way. Importantly, it is the basic-level categories (e.g., “bird,” “dog”) that are prioritized in human behavior^58,59^. Future studies might explore fine-tuning on more ecologically aligned, basic-level datasets (e.g., Ecoset^60^) to encourage a similar cue-reliance profile as humans.

A similar explanation may also apply to the same tendency shown by zsCLIP in the **Fil** condition (high performance but low error consistency with humans). zsCLIP is not trained on ImageNet, but its training data typically consist of image-caption pairs assembled from the internet, which may contain more detailed descriptions than humans would extract from briefly presented images alone. Taken together, these findings suggest that to enhance human-alignment, the granularity of the information used during training should be reconsidered so that it better matches how humans naturally perceive visual objects. We expect that CLIP models trained and fine-tuned on such a training diet may exhibit behavior that is more closely aligned with human visual recognition.

More broadly, an important insight from our Experiment 1 is that depending on whether the global shape cue is present, classification accuracy either predicts error consistency with humans or is decoupled from it. This suggests that the common objective of maximizing task performance alone could make DNNs recognize the images that are challenging for humans, and not necessarily lead to a globally human-like model. To achieve a truly human-like model that succeeds when humans succeed and fails when humans fail, training objectives that explicitly incorporate human behavioral data may be necessary^61^. By contrast, in shape-present conditions, human performance remains the near upper bound for today’s machine performance.

In addition, our results suggest that object recognition in both humans and DNNs often depends on the interaction of multiple visual cues. For example, human accuracy in the **Fil** condition (70.8%; local parts + texture) was higher than would be expected from combining performance in the **Tex** condition (texture alone; 18.4%) and the **LinFil** condition (local parts alone; 26.9%), and higher than the **Sil** condition (global shape alone; 58.4%). This suggests that the joint presence of both types of local information—texture and local parts—appears to be especially beneficial to object recognition. Indeed, the combination of texture and local parts information might form high-level texture patterns that are critical to human perception of materials^62^. Therefore, the low accuracies in the **Tex** and **LinFil** conditions should not be taken to mean that texture or local-part information is uninformative, which aligns with prior work demonstrating the substantial impact of texture- and local-part cues on visual cortical activity^23,63,64^. Examining such cue-interaction patterns in greater detail across humans and DNNs may provide further insight into the mechanisms underlying object recognition, and remains an important topic for future work.

In Experiment 2, no DNNs—including ftCLIPs that excelled in the **Sil** condition of Experiment 1—attained human-level accuracy when global shape was decoupled from local shape. It might be argued that this failure stems from the absence of **Sparse** or **Dense**-like images in the DNNs’ training data; however, human observers likewise lack explicit experience with such stimuli yet readily perceive their global form. This contrast suggests that human vision harnesses strong inductive biases that automatically favor the global shape over local details. A truly human-aligned model should similarly generalize to such fragmented silhouettes without direct exposure during training. This finding thus highlights the divergence between machine vision and human vision, especially in their weighting of local versus global cues. Intriguingly, we found that DNNs do not entirely lose the global-shape information; rather, these signals are masked by dominant local-feature-driven activations. Indeed, our ablation experiment was striking in showing that when ∼30–80% of units in the penultimate layer, which were tuned to local structure, were set to 0, the DNN’s performance often improved when perceiving these fragmented silhouettes.

Currently, the biological underpinnings of the strong inductive bias toward the global shape in human vision remain unclear. Several possibilities exist: First, unlike DNNs, the ventral stream might inherently encode global silhouettes more strongly than local shape, although this possibility is cast into doubt by recent fMRI evidence^23^. Alternatively, regions downstream to the ventral visual pathway, the prefrontal cortex, might selectively upweight global-shape signals when guiding behavior^23^. Finally, the global shape might be processed in regions outside the ventral stream, such as the dorsal visual stream^49,65^. Additional empirical investigation will be necessary to unravel the neural mechanisms underlying human object recognition’s strong reliance on the global shape. Of note, a recent study suggests that this strong global shape bias of human object perception might not be matched by macaque monkeys^66^, underscoring the importance of mechanistic investigation in humans. Finally, integrating such empirical insights into future DNN models will be an essential next step toward constructing artificial agents that truly mirror human object recognition.

In conclusion, we reveal systematic differences in how DNNs and humans leverage visual cues to accomplish object recognition. We found that training on manipulated images, incorporating recurrent connections, and improving alignment to ventral stream neural recordings so far did not produce human-aligned object recognition behavior in DNNs. Our findings illuminate when performance is a good guide for human alignment and when it is not, and underscore the importance of understanding the strong inductive bias of human vision to rely on the global shape cue. Overall, we found that the training approach had a larger impact on a model’s behavioral alignment with humans than the model architecture. Together, these results shed new light on how the human brain accomplishes object recognition with limited cues and point to concrete avenues for further enhancing the human alignment of machine vision.

### Limitations of the study

Our results indicate that CLIP models are often more human-aligned than other models, and we speculate that natural-language supervision is one crucial factor underlying this high human alignment. However, because CLIPs differ from models trained solely on ImageNet in several respects (e.g., training dataset size: ImageNet ∼1.2 M images vs. CLIP 400 M–2 B images, and dataset diversity), further work is needed to pinpoint which factors drive their human-like behavior.

In addition, although we extensively investigated human-DNN alignment at the level of output behavior (i.e., object labels), high output alignment does not necessarily guarantee that humans and DNNs process image information in the same way internally. Such internal alignment is also essential for a model to serve as a strong computational account of human vision. As noted above, because neural recordings for this image set are not currently available, we cannot yet convincingly assess this form of internal alignment.

## RESOURCE AVAILABILITY

### Lead contact

Requests for further information and resources should be directed to and will be fulfilled by the lead contact, Biyu He.

### Materials availability

This study did not generate new unique reagents.

### Data and code availability

- Data including human psychophysics and DNNs’ predictions can be found at: https://github.com/BiyuHeLab/DNNObjectRec_Kato2026 (DOI: <u>10.5281/zenodo.21227163</u>). The images used in the experiments are not publicly available due to copyright restrictions of the ImageNet database, but the above GitHub repository includes ImageNet image IDs and code to generate the images for each condition, allowing the recreation of our image set.
- The code supporting this work is available in an open GitHub repository: https://github.com/BiyuHeLab/DNNObjectRec_Kato2026 (DOI: <u>10.5281/zenodo.21227163</u>).
- All other data reported in the manuscript will be shared by the Lead contact upon request.

## ACKNOWLEDGEMENTS

We thank Eric K. Oermann for illuminating discussions and Animesh Mishra for technical assistance. This research was supported by a W.M. Keck Foundation grant (to B.J.H.), a grant from National Institutes of Health (R01EY032085, to B.J.H.) and fellowship awards from Astellas Foundation and Uehara Memorial Foundation (to M.K.).

## AUTHOUR CONTRIBUTIONS

M.K. performed all the experiments and analyses. M.K. and B.J.H designed research and co-wrote the manuscript.

## STAR★METHODS

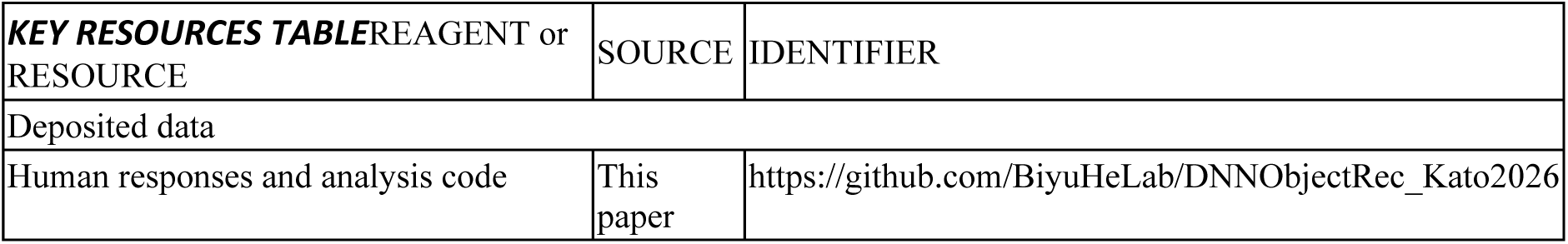

### EXPERIMENTAL MODEL AND STUDY PARTICIPANT DETAILS

In Experiment 1, 101 subjects were recruited from the Prolific.co subject pool. Among them, 18 subjects were excluded at the initial stage because they failed to meet the inclusion criteria: using a desktop monitor, having normal or corrected-to-normal vision, not being color blind, and passing the practice session of the experimental task (see *Experimental procedure* for details). Twenty-one subjects were excluded because they did not complete the study. Two additional subjects were excluded from analysis: one did not follow instructions (often providing multiple answers in a single trial), and the other failed to respond in many trials. Consequently, data from 60 subjects (27 female, 2 unreported; mean age 38.93, range 19–73) were included in the analysis.

In Experiment 2, 21 subjects were recruited from the Prolific.co subject pool. Among them, 5 subjects were excluded at the initial stage because they did not meet the inclusion criteria, which were the same as those in Experiment 1. Six subjects were excluded because they did not complete the study. Consequently, data from 10 subjects (1 female, 2 unreported; mean age 39.70, range 26–61) were included in the analysis.

All subjects provided written informed consent, and the experiment was approved by the Institutional Review Board of New York University School of Medicine (protocol #S21-00218).

Because the study focused on the recognition of common objects that were expected to be broadly familiar across participants, regardless of gender or other demographic variables, we did not have any gender-related hypotheses. Accordingly, we did not analyze the influence of gender or collect demographic variables beyond age and gender. Participants were randomly assigned to subject groups without considering demographic variables (see Experimental procedure).

### METHOD DETAILS

#### Experiment 1

##### Stimulus selection

All stimuli were drawn from the validation set of the ImageNet Large Scale Visual Recognition Challenge 2012 database, which comprises 50 images for each of 1,000 classes^67^. To ensure broad coverage of everyday objects and animals, we defined 12 basic categories within each of four coarse groups—mammals, non-mammal animals, man-made objects, and natural objects—resulting in 48 basic categories in total (**Fig. S1A**). Using the WordNet lexical hierarchy^35^, we mapped each ImageNet class onto a basic category: an ImageNet class was assigned to a basic category if its name was identical to, a synonym of, or a hyponym of the basic category name. To better reflect human conceptual grouping, we additionally made a small number of manual assignments for classes not captured by the automated criteria (i.e., “hare” to rabbit, “cup” to mug, “ear” to corn, and “agaric”, “bolete”, “coral fungus”, “earthstar”, “gyromitra”, “hen-of-the-woods”, and “stinkhorn” to mushroom). Within each basic category, we then randomly selected five exemplar images from the validation set, discarding any images that lacked a clear object-background boundary, depicted a nonreal object (e.g., a stuffed doll), contained overlapping objects of different categories, or measured less than 256×256 pixels. This procedure yielded 240 exemplars in total.

##### Image manipulation

First, each image was cropped and resized to 256×256 pixels, and its background was manually removed in Adobe Photoshop to create original images (“**Ori**”). For each Ori image, we generated 6 manipulated conditions (**Figs. 1A, S1B, S2**), each of which disrupts one (**Fil, Lin, SilTex**) or two (**Sil, Tex, LinFil**) of the three cue types: global shape, texture, and internal parts. Although terms such as “global shape’’ and “texture’’ are commonly used in the field, there is no single agreed-upon definition. Therefore, below we clarify how we use these terms, how our definitions relate to previous work, and how we disrupted each cue.

We define global shape as the object’s overall outline contour, which can be perfectly isolated from other cues by a binary object mask (i.e., silhouette, “**Sil”**). This definition matches prior studies that evaluate object recognition from shape alone^20,21,25,68^. To disrupt global shape while preserving the other two cues, we generated filled (**“Fil”**) images by repeatedly duplicating and shifting the original image until the entire frame was covered, producing a crowded presentation that disrupts the overall contour but preserves local shapes and texture (**Figs. 1A, S1B**). We note that a previous study labeled a similar image condition “texture images”^20^; however, we consider these images to contain both internal-part information (i.e., local shape information) and texture.

To extract texture, we use the VGG19-based texture-synthesis algorithm^69^. Each convolutional or pooling layer in VGG19 produces multiple 2D feature maps, each of which functions as a distinct nonlinear filter bank that extracts specific visual features; spatial locations on these maps correspond to positions in the input image. Here, the inner products between all pairs of feature maps, or the Gram matrix, encode the presence of individual features and their co-occurrence while discarding spatial location information, which matches the usual notion of texture as location-invariant, repeated visual features. Indeed, the original paper demonstrated that an image composed of spatially uniform texture can be reproduced by iteratively generating a new image that matches the Gram matrix of the original image^69^. When this synthesis procedure is applied to natural images, it produces a spatially scrambled (“texturized”) version of the original image. Consequently, recent studies commonly use this method to extract texture information from natural images^20,22–24,70^. Note that a more lenient definition of texture has also been used in some studies, in which images with cluttering of the same object or by cropping within an object’s contour are treated as texture images^20,21^; however, as mentioned above, such images have the confounding cue of local shape information that remains intact.

Importantly, the complexity of texture-like features preserved in a synthesized image is controlled by the choice of CNN layers used to compute the Gram matrices (**Fig. S1D**)^69^. When only early layers are included, the synthesized image is highly scrambled and no human-recognizable local parts remain. As progressively higher layers are included, spatial scrambling is reduced and increasingly recognizable local structure is preserved. Although prior studies typically include a broad range of layers (from early layers up to layer 4 or 5 out of 5 layers in total) to extract texture information, the resulting images often contain elements that are not strictly texture, such as individual facial parts^22–24^. To address this issue and to better separate texture from local shape information in our stimuli set, we first determined which layer choice best captures texture in a stricter sense for our stimuli. To avoid influence from the gray background, we applied the texture synthesis procedure to the **Fil** images and evaluated five layer choices, including all layers up to pooling layers 1-5 (**Fig. S1D**). Based on visual inspection, individual local parts (e.g., facial parts, car headlights/wheels) become recognizable when layer 3 or higher layers are included, whereas using layers only up to layer 2 preserves texture while removing such distinctive local shape cue. Thus, in this study, we generate images preserving only texture information (**“Tex”** images) by matching the Gram matrix computed over layers up to pooling layer 2 (2,000 iterations; all other settings at default^69^). In addition, combining **Tex** images with silhouette masks yielded “**SilTex**” images, which contain both global shape and texture cues.

Our **Tex** images created using the above procedure isolate texture information. Conversely, to remove the texture information and leave intact both the global shape cue and internal parts information (i.e., local shape cues), we used an automatic line-drawing image generation algorithm^71^. The resulting line drawings reliably depict internal parts as well as contour outlines in black and white, thereby minimizing texture information (**Fig. S1B**)^72^. To ensure a clear global shape, we superimposed each raw line drawing onto the corresponding silhouette image to generate the **Lin** image, which preserves both global shape and internal parts (**Figs. 1A, S1B, S2**). Finally, to disrupt the global shape and isolate only internal-parts information, we applied the same duplicating and shifting procedure used for **Fil** images to the raw line drawings, producing the line-filled (**“LinFil”**) stimuli.

##### Experimental procedure

The experiment was implemented on Gorilla.sc, with each subject using their own desktop monitor. Screen sizes were calibrated individually using a credit card, and subjects were instructed to maintain a consistent viewing distance. Before beginning the main task, each subject completed practice trials under timed and untimed conditions.

Because DNNs do not have continual learning at test time (i.e. weights are frozen at test time), whereas human subjects continuously learn throughout the task and can use past images from one condition to disambiguate related images in a different condition^73^, we implemented strict controls to mitigate learning effects in human subjects to ensure a fair comparison between human responses and DNNs. This was done using the following strategy.

The main session lasted approximately 60 minutes on average and comprised 4 consecutive blocks. The first two blocks presented the six manipulated conditions (**Fil, Lin, LinFil, Tex, Sil, SilTex**) in a random order, and the final two blocks presented the original images (**Ori**). The original images were presented last to prevent subjects from learning from prior exposure to the intact images, which could make otherwise unrecognizable manipulated images recognizable in later trials. All 240 exemplars appeared in the **Ori** blocks, whereas for each manipulated condition, a given subject saw a different subset of 40 exemplar images (**Fig. S1C**, top). Thus, for each exemplar image, a given subject saw it twice: once in a specific (randomly assigned) manipulated condition, and once as the original image. This design thus avoided potential learning effects arising from repeated presentations of the same exemplar image across different manipulation conditions (e.g., seeing the blue jay in **SilTex** might facilitate recognizing the same exemplar in **Tex**; see examples in **Fig. S2**).

Specifically, the 240 exemplars were partitioned into six subsets of 40 images, with each subset including 1 exemplar from 40 different basic categories, balanced across the four coarse category groups (**Fig. S1C**, top). The 60 subjects with successful data collection formed 10 groups of 6 subjects each. Within each group, which subset of 40 images was assigned to which manipulation was counterbalanced across subjects (**Fig. S1C**, bottom), such that each subject saw a given exemplar image only in a single manipulation condition and, collectively, the six subjects within a group saw all 240 exemplars in each manipulated condition and no two subjects within a group saw the exact same manipulated image. New partitions were generated for each group of six subjects by shuffling the position of the 5 exemplars within each basic category (i.e., each column) in the matrix shown in **Fig. S1C**, top.

Within each block, trials were presented in a pseudorandom order and included 120 main trials and 24 dummy trials (explained below). Main trials began with a fixation cross on a gray background for an interval of 1,300–1,700 ms (uniformly distributed), followed by a stimulus image presented at central fixation (subtending 10° visual angle) for 500 ms (**Fig. 1B**). After image offset, a text input box appeared and remained until the subject pressed Enter or until 6 seconds elapsed. Subject were instructed to type the object or animal name that first came to mind, using specific exemplar labels (e.g., “cat”) rather than broad categories (e.g., “animal”), followed by pressing Enter. Dummy trials appeared unpredictably between main trials to ensure that subjects maintained fixation: during a dummy trial, an image outside the selected 48 basic categories was shown, and at a random time between 1,000 ms before image offset and image offset, the fixation cross briefly changed color for 100 ms. Subjects were instructed to press the space bar as soon as they detected this change. No text input box appeared in dummy trials. During practice, subjects who failed to respond to the fixation-color change within 1,000 ms after image offset on two or more of five dummy trials repeated the practice. Subjects who did not meet this threshold on a second attempt were excluded.

##### Evaluation of human responses

Typed responses were extracted from Gorilla.sc and processed in MATLAB (R2024a) with custom scripts. Each response was compared automatically to answer keys defined via WordNet^35^: any response that was a hyponym or synonym of the correct basic category, or that exactly matched the category name, was marked correct. The ImageNet class names we manually added were also marked as correct (see *Experimental Stimuli*). Responses with minor spelling errors were manually reviewed and corrected when appropriate.

##### Evaluation of model responses

We evaluated over 200 DNNs on the identical set of 240 exemplars × 7 conditions used with human observers. Eleven publicly available models were primarily analyzed (**Fig. 1C**). Four were ImageNet-trained models from the PyTorch/Torchvision model zoo^74^: ResNet50 (inResNet); ConvNeXt_Large (inConvNeXt)^75^; CORnet-S (inCORnet)^37^; and ViT_L_16 (inViT). Three had been trained on blurred or stylized versions of ImageNet: ResNet50_Strong-blur (blResNet)^32^, CORnetS_Strong-blur (blCORnet)^32^, resnet50_trained_on_SIN (stResNet)^20^. Two were zero-shot CLIP models trained on LAION2B^76^ available in OpenCLIP^77^: CLIP-convnext_large_d.laion2B-s26B-b102K-augreg (zsConvNeXt), CLIP-ViT-L-14-laion2B-s32B-b82K (zsViT). The last two were CLIP models fine-tuned on ImageNet classification available in Pytorch-Image-Models (timm) library^78^: convnext_large_mlp.clip_laion2b_augreg_ft_in1k_384 (ftConvNeXt), vit_large_patch14_clip_224.laion2b_ft_in12k_in1k (ftViT). For models other than zsCLIPs, each image was passed through the network and the top 1 predicted class was recorded. For zsCLIP models, we computed cosine similarity between the image embedding and text embeddings for “an image of a [class name]” across all 1,000 ImageNet classes, selecting the class with the highest similarity. The Stable Diffusion classifier was implemented as described in ^79^. A model response was deemed correct if the predicted class fell within the correct basic category, as defined in the *Experimental Stimuli* section. In a broader analysis, we applied the same evaluation to all models available in the Brain-Score database^44^, excluding redundant models and those with **Ori** accuracy below 50% (mostly untrained models). This added 198 models to our set of 11 (Table S1).

##### Recognition accuracy

For human subjects and each DNN, recognition accuracy in the **Ori** condition was calculated as the percentage of correctly recognized exemplars out of 240. In manipulated conditions, exemplars not recognized in the **Ori** condition by a given subject or DNN model were excluded and accuracy was computed as the percentage correct out of the remaining exemplars. This approach isolates the impact of image manipulations independent of baseline recognition ability. For manipulated conditions in humans, data from each group of six subjects were pooled to form a “pooled subject,” who completed the full image coverage across those six individuals (**Fig. S1C**). We note that pooling data across multiple human subjects to compare with DNNs is a standard approach^11,80,81^, due to both the time limitation of human subjects and their learning and fatigue effects (DNNs, by contrast, generally do not learn at test time).

To compare model accuracy against human performance, we computed, for each pooled human subject, the difference between that subject’s accuracy and the model’s accuracy. A two-sided Wilcoxon signed-rank test assessed whether the average of these differences significantly deviated from zero. P-values were Bonferroni-corrected across the number of models tested.

##### Error consistency

Error consistency was used to quantify whether two conditions or two observers agree (both correct or both incorrect) on more images than would be expected if their responses were independent given their accuracies^11,36^. This measure, equivalent to Cohen’s κ, is defined for observers *i* and *j* by

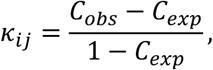

where *CC_obs_*is the proportion of images with matching responses, and

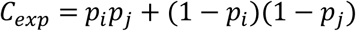

is the expected agreement under independence, with *p_i_*and *p_i_* denoting the accuracies of observers *i* and *j*.

Within-subject, between-condition error consistency (**Fig. 1E**) was computed for every pair of image conditions, separately for each pooled human subject and for each DNN. Only exemplars that were correctly recognized in the **Ori** condition were included. To determine whether the observed error consistency between conditions exceeded chance, we performed an image-label permutation test for each observer and each condition pair. Specifically, we generated a null distribution by randomly shuffling the mapping between images and responses (correct/incorrect) in one condition and recomputing error consistency. After 10,000 permutations, the one-sided p-value was defined as the proportion of permutations in which the permuted error consistency exceeded the observed value. For the human data, we also performed a group-level statistical test: a one-sided Wilcoxon signed-rank test against zero on error consistency was conducted across the 10 pooled subjects. P-values were FDR-corrected across all condition pairs.

Between-observer error consistency (**Figs. 2-4, S3**) was computed between each DNN and every pooled human subject, as well as between all pairs of pooled subjects. Only exemplars correctly recognized by both observers in the **Ori** condition were included in each pairwise calculation. To determine whether a DNN’s error pattern deviates from the range of human variability, we employed a bootstrap procedure to generate a null distribution under the assumption that the model’s responses are exchangeable with human responses. For each DNN and condition, we conducted 2,000 bootstrap iterations. In each iteration, nine human pooled-subject response vectors were sampled with replacement and the DNN’s response vector was appended to this set to form a “pseudo-human” group of ten observers. In addition, one of the original ten pooled-subject response vectors was randomly selected to serve as the “pseudo-DNN”. Cohen’s κ was computed for all pseudo-human to pseudo-human pairs and for each pseudo-human to pseudo-DNN pair. The difference between the mean pseudo-human to pseudo-human κ and the mean pseudo-human to pseudo-DNN κ in each iteration constituted the bootstrap statistic under the null hypothesis that the DNN’s error pattern is indistinguishable from human variability. Because resampling can include the same observer multiple times—yielding self-comparisons with κ = 1 (the highest value) that rarely occur in real data—we excluded any such self-pairs before averaging. We calculated the observed difference from the actual human and DNN responses in the same manner, and defined the one-sided *p*-value as the proportion of bootstrap iterations in which the null difference exceeded the observed difference. P-values were Bonferroni-corrected across models within each condition.

##### Correlation between IT predictability and behavioral similarities to humans

IT predictability scores were obtained from the Brain-Score database^44,45^, and only models with available IT scores (128 out of 209) were included. Human-DNN behavioral similarity in our data set was quantified in three ways for each image condition:

1. The absolute difference in recognition accuracy between the model and the mean of pooled human subjects: 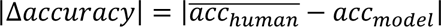 (**Fig. S4A**);
2. The difference between human-to-human error consistency and human-to-DNN error consistency: 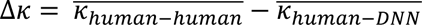 (**Fig. S4B**); and
3. a combined metric defined as the root-sum-square of the normalized differences in accuracy and consistency (**Fig. S4C**), as described below. We normalized both model and human measures using z-scores computed across the full set of 128 models. Concretely, for accuracy we computed a z-score for each model *m* and for the human average as

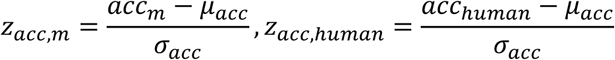

where *acc_m_* is model *m*’s accuracy, *acc*_human_ is the average human accuracy for that condition, and *μμ_acc_*and *σσ_acc_*are the mean and standard deviation of accuracies calculated across the 128 models. The same procedure was applied to error-consistency values:

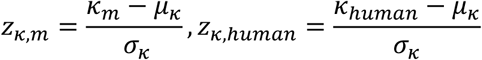

where *κ_m_*is average of model *m*’s error consistency with humans, *κκ*_human_ is the average human-to-human error consistency for that condition, and *μ_k_*and *σ_k_*are the mean and standard deviation of error consistencies calculated across the 128 models. We combined normalized accuracy and consistency into a single distance from human performance by taking the root-sum-square of the differences between model and human z-scores:

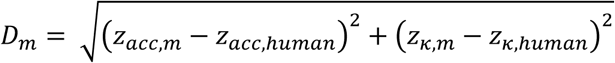

*D_m_*represents the Euclidean distance in the two-dimensional (normalized accuracy, normalized error consistency) space between model *m* and the human average for that condition. Finally, for each condition, we computed the Pearson correlation between IT predictability and each of these three behavioral-similarity measures (magenta annotations in **Fig. S4**).

##### Correlation between behavioral similarities to humans measured by our image set and the Brain-Score behavioral benchmark

As an overall measure of DNNs’ behavioral similarity to humans in the Brain-Score, we used the average score across 48 behavioral benchmarks (labeled “behavior_vision” in Brain-Score) available as of April 2026. For each condition, we computed the Pearson correlation between this mean score and each of the three behavioral-similarity measures from our image set described above (i.e., |Δaccuracy|, Δ*κ*, and *D_m_*; blue annotations in **Fig. S4**).

##### Exemplar-wise representational similarity analysis

To investigate how different training approaches influence the internal representations of visual features in DNNs, we conducted an exemplar-wise representational similarity analysis (RSA)⁵⁷. We selected four representative layers spanning the hierarchy of processing within each model: the first activation layer, the middle activation layer, the final activation layer, and the penultimate layer (**Fig. 5A**; left; see Table S3 for layer labels). For each of the 240 exemplars, we computed a representational dissimilarity matrix (RDM) at each layer, using the correlation distance (1 – Pearson’s *r*) between activation patterns evoked by the seven image variants derived from the same exemplar (**Fig. S6A**; left). We also constructed three model RDMs reflecting hypothesized dissimilarity structures based on shape, texture, and internal-parts information (**Fig. S6A**; right). In each model RDM, image pairs that both retained the visual feature of interest were assigned a dissimilarity value of 0, while all other pairs were assigned a value of 1. We quantified the alignment between each layer RDM and each model RDM using Spearman’s rank correlation. The resulting correlation coefficients were transformed using Fisher’s *z*-transformation and averaged across exemplars. We used one-sided one-sample *t*-tests to assess whether the mean correlation significantly exceeded zero, and two-sided paired *t*-tests to compare correlations across models sharing the same architecture but differing in training. P values were Bonferroni-corrected across layers.

##### Leave-one-exemplar-out object decoding

To evaluate the amount of object information encoded in each hidden layer, we performed a leave-one-exemplar-out object decoding analysis (**Fig. 5A**). For each image condition and layer, we computed Pearson’s correlation between the activation pattern of a left-out exemplar and all remaining 239 exemplars. The resulting correlation values were then averaged within each of the 48 basic-level object categories, and the category with the highest correlation value was the output label. If this output label corresponded to the actual category of the left-out exemplar, the exemplar was considered correctly decoded. This process was repeated such that each exemplar was left out once, and decoding accuracy was calculated as the percentage of correctly predicted exemplars.

To determine statistical significance, we conducted a label permutation test. Specifically, we randomly shuffled the category labels and repeated the decoding procedure 1,000 times to generate a null distribution of decoding accuracies. The one-sided P value was defined as the proportion of permutations in which the shuffled accuracy exceeded the observed decoding accuracy. All P values were Bonferroni-corrected across layers.

#### Experiment 2

##### Experimental Stimuli

To generate silhouette images where global shape is decoupled from local shape, we created a variant of **Sil** images by filling object regions with randomly placed cross-shaped elements. Since silhouette images that most humans were unable to recognize are not worth testing in this experiment, we only included 171 out of 240 exemplars for which human accuracy in the **Sil** condition exceeded 30% in Experiment 1. Starting with a set of Sil images (256×256 pixels), we randomly distributed cross-shaped symbols (i.e., “+” marks) within the boundaries of each object silhouette. The density and size of these crosses were parametrically controlled: cross diameter, 10 pixels; placed crosses at a density of 0.03 and 0.075 per silhouette pixel for **Sparse** and **Dense** conditions, respectively. The total number of crosses for each image was computed as the product of the total silhouette area and the density. Half-sized **Sparse** and **Dense** images were generated by downscaling the original images by a factor of two (128 × 128 pixels) and embedding them within a gray margin so that the overall image size remained 256 × 256 pixels (**Fig. S5B**).

##### Experimental procedure

The experiment was implemented on Gorilla.sc, with each subject using their own desktop monitor. Screen sizes were calibrated individually using a credit card, and subjects were instructed to maintain a consistent viewing distance. Subjects were instructed to type the object or animal name that first came to mind, using specific exemplar labels (e.g., “cat”) rather than broad categories (e.g., “animal”). Before beginning the main task, each subject completed practice trials under timed and untimed conditions.

The main session lasted approximately 50 minutes on average and comprised 6 consecutive blocks. The first 2, middle 2, and the last 2 blocks presented the **Sparse, Dense**, and **Sil** conditions, respectively. The conditions were ordered from most challenging to least challenging to prevent recognizing an easier image disambiguating the corresponding harder image. All 171 exemplars appeared in each condition in a pseudorandom order. Each block consisted of 85 or 86 main trials and 17 dummy trials. The timeline of these trials was exactly the same as that of Experiment 1 (**Fig. 1B**).

##### Evaluation of human and model responses

In the analysis where DNNs’ top-5 response was examined (**Fig. S5A**), a response was counted as correct if the correct label appeared among the top five predictions, instead of the top one. Otherwise, these procedures were performed in the same manner as in Experiment 1.

##### Recognition accuracy and Error consistency

Because every subject in Experiment 2 viewed the full stimulus set, we analyzed each subject’s data individually rather than pooling across subjects. Otherwise, these analyses were conducted following the same procedures as in Experiment 1.

##### Correlation between IT predictability and behavioral similarities to humans

Performed in the same manner as in Experiment 1.

##### Leave-one-exemplar-out decoding

For this analysis, we required at least two exemplars per basic category. 41 out of the 48 categories met this criterion (due to Experiment 2 having a total of 171 exemplar images; see *Experimental Stimuli* above) and were used in the decoding analysis, yielding a chance-level decoding accuracy of 1/41. All other procedures mirrored those of Experiment 1.

##### Silencing locally tuned units

To determine whether object information defined by global shape remains encoded in hidden layers despite dominant local-feature activations, we evaluated recognition performance after silencing units that respond preferentially to local cross patterns regardless of global silhouettes. For each unit in the penultimate layer, we tested whether its activation to the **Sparse** or **Dense** images differed significantly from its activation to the seven conditions of those exemplars used in Experiment 1. Specifically, we performed a two-sided two-sample *t*-test for each unit, comparing its responses to the **Sparse** or **Dense** set (240 images) against responses to the other image conditions (240×7 =1680 images). Units exhibiting *p*<0.05 after Bonferroni correction (across all penultimate-layer units) were classified as locally tuned (**Fig. 5C**; left). Note that this procedure can detect units that show either greater or lesser activation in the **Sparse/Dense** conditions compared with the other seven conditions. We considered units that were either activated or suppressed in the **Sparse/Dense** conditions to be locally tuned, because unit activations are multiplied by downstream weights that can be positive or negative. Thus, suppression of a unit can also contribute to an increased probability for objects with cross-like features. We then assessed each DNN’s recognition accuracy after setting these units’ activations to zero and passing the modified activations through the final classification layer (**Fig. 5C**; right).

zsCLIPs were excluded from this analysis because their predictions additionally rely on a separate text-embedding space. In standard networks with a fully connected classification layer, the activation of each unit in the penultimate layer is multiplied by learned weights and passed to the next layer; setting a unit’s activation to zero therefore removes that unit’s contribution to the downstream class scores. By contrast, in zsCLIP models, predictions are based on the similarity between the overall pattern of visual embeddings and text embeddings. Because the model evaluates embedding vectors as whole patterns, setting individual unit activations in the visual embeddings to zero does not simply eliminate that unit’s influence, making the result not straightforwardly interpretable. In contrast, CLIP models that were fine-tuned on ImageNet were included since they have an additional fully connected classification layer, which allows them to behave like standard networks without relying on text embeddings.

### QUANTIFICATION AND STATISTICAL ANALYSIS

All statistical tests were performed using MATLAB functions (Statistics and Machine Learning Toolbox, R2021B) or custom-written MATLAB code. For each analysis, the statistical test, the number of observations, multiple-comparison correction, and alpha level are reported in the corresponding figure legend and *Method details*, and full statistical details are provided in the supplementary Excel file.

## ADDITIONAL RESOURCES

We intend to submit the entire image set and human behavioral responses to Brain-Score as a behavioral benchmark (https://www.brain-score.org/). This will allow anyone to evaluate a model’s outputs on our image set and directly compare them with human responses.

## SUPPLEMENTAL ITEMS

**DataS1:** Raw statistical data related to Figures 1–6 and S3, S5–S7.

## REFERENCES

1. Kar, K., and DiCarlo, J.J. (2024). The quest for an integrated set of neural mechanisms underlying object recognition in primates. Annual Review of Vision Science 10.

2. Alzubaidi, L., Zhang, J., Humaidi, A.J., Al-Dujaili, A., Duan, Y., Al-Shamma, O., Santamaría, J., Fadhel, M.A., Al-Amidie, M., and Farhan, L. (2021). Review of deep learning: concepts, CNN architectures, challenges, applications, future directions. Journal of Big Data 8, 53. 10.1186/s40537-021-00444-8.

3. He, K., Zhang, X., Ren, S., and Sun, J. (2015). Delving deep into rectifiers: Surpassing human-level performance on imagenet classification. pp. 1026–1034.

4. Shankar, V., Roelofs, R., Mania, H., Fang, A., Recht, B., and Schmidt, L. (2020). Evaluating Machine Accuracy on ImageNet. In D. Hal, III, and S. Aarti, eds. Proceedings of the 37th International Conference on Machine Learning. PMLR.

5. Cichy, R.M., Khosla, A., Pantazis, D., Torralba, A., and Oliva, A. (2016). Comparison of deep neural networks to spatio-temporal cortical dynamics of human visual object recognition reveals hierarchical correspondence. Scientific Reports 6, 27755. 10.1038/srep27755.

6. Güçlü, U., and van Gerven, M.A. (2015). Deep Neural Networks Reveal a Gradient in the Complexity of Neural Representations across the Ventral Stream. J Neurosci 35, 10005–10014. 10.1523/jneurosci.5023-14.2015.

7. Horikawa, T., and Kamitani, Y. (2017). Generic decoding of seen and imagined objects using hierarchical visual features. Nat Commun 8, 15037. 10.1038/ncomms15037.

8. Yamins, D.L., Hong, H., Cadieu, C.F., Solomon, E.A., Seibert, D., and DiCarlo, J.J. (2014). Performance-optimized hierarchical models predict neural responses in higher visual cortex. Proc Natl Acad Sci U S A 111, 8619–8624. 10.1073/pnas.1403112111.

9. Yamins, D.L.K., and DiCarlo, J.J. (2016). Using goal-driven deep learning models to understand sensory cortex. Nature Neuroscience 19, 356–365. 10.1038/nn.4244.

10. Conwell, C., Prince, J.S., Kay, K.N., Alvarez, G.A., and Konkle, T. (2024). A large-scale examination of inductive biases shaping high-level visual representation in brains and machines. Nat Commun 15, 9383. 10.1038/s41467-024-53147-y.

11. Geirhos, R., Narayanappa, K., Mitzkus, B., Thieringer, T., Bethge, M., Wichmann, F.A., and Brendel, W. (2021). Partial success in closing the gap between human and machine vision. Advances in Neural Information Processing Systems 34, 23885–23899.

12. Szegedy, C., Zaremba, W., Sutskever, I., Bruna, J., Erhan, D., Goodfellow, I., and Fergus, R. (2013). Intriguing properties of neural networks. arXiv preprint arXiv:1312.6199.

13. Geirhos, R., Temme, C.R., Rauber, J., Schütt, H.H., Bethge, M., and Wichmann, F.A. (2018). Generalisation in humans and deep neural networks. Advances in neural information processing systems 31.

14. Dodge, S., and Karam, L. (2017). A study and comparison of human and deep learning recognition performance under visual distortions. 2017 26th international conference on computer communication and networks (ICCCN), 1–7.

15. Jang, H., McCormack, D., and Tong, F. (2021). Noise-trained deep neural networks effectively predict human vision and its neural responses to challenging images. PLoS biology 19, e3001418. 10.1371/journal.pbio.3001418.

16. Jang, H., and Tong, F. (2021). Convolutional neural networks trained with a developmental sequence of blurry to clear images reveal core differences between face and object processing. Journal of vision 21, 6. 10.1167/jov.21.12.6.

17. Muttenthaler, L., Dippel, J., Linhardt, L., Vandermeulen, R.A., and Kornblith, S. (2022). Human alignment of neural network representations. arXiv preprint arXiv:2211.01201.

18. Nonaka, S., Majima, K., Aoki, S.C., and Kamitani, Y. (2021). Brain hierarchy score: Which deep neural networks are hierarchically brain-like? iScience 24, 103013. 10.1016/j.isci.2021.103013.

19. Gokce, A., and Schrimpf, M. (2024). Scaling laws for task-optimized models of the primate visual ventral stream. arXiv preprint arXiv:2411.05712.

20. Geirhos, R., Rubisch, P., Michaelis, C., Bethge, M., Wichmann, F.A., and Brendel, W. (2018). ImageNet-trained CNNs are biased towards texture; increasing shape bias improves accuracy and robustness. International conference on learning representations.

21. Baker, N., Lu, H., Erlikhman, G., and Kellman, P.J. (2018). Deep convolutional networks do not classify based on global object shape. PLoS Comput Biol 14, e1006613. 10.1371/journal.pcbi.1006613.

22. Gatys, L.A., Ecker, A.S., and Bethge, M. (2017). Texture and art with deep neural networks. Current Opinion in Neurobiology 46, 178–186. 10.1016/j.conb.2017.08.019.

23. Jagadeesh, A.V., and Gardner, J.L. (2022). Texture-like representation of objects in human visual cortex. Proc Natl Acad Sci U S A 119, e2115302119. 10.1073/pnas.2115302119.

24. Brendel, W., and Bethge, M. (2019). Approximating cnns with bag-of-local-features models works surprisingly well on imagenet. arXiv preprint arXiv:1904.00760.

25. Baker, N., and Elder, J.H. (2022). Deep learning models fail to capture the configural nature of human shape perception. iScience 25, 104913. 10.1016/j.isci.2022.104913.

26. Jacob, G., Pramod, R.T., Katti, H., and Arun, S.P. (2021). Qualitative similarities and differences in visual object representations between brains and deep networks. Nat Commun 12, 1872. 10.1038/s41467-021-22078-3.

27. Baker, N., Lu, H., Erlikhman, G., and Kellman, P.J. (2020). Local features and global shape information in object classification by deep convolutional neural networks. Vision Research 172, 46–61. 10.1016/j.visres.2020.04.003.

28. Lonnqvist, B., Scialom, E., Gokce, A., Merchant, Z., Herzog, M.H., and Schrimpf, M. (2025). Contour Integration Underlies Human-Like Vision. arXiv preprint arXiv:2504.05253.

29. Jaini, P., Clark, K., and Geirhos, R. (2023). Intriguing properties of generative classifiers. arXiv preprint arXiv:2309.16779.

30. Wang, A.Y., Kay, K., Naselaris, T., Tarr, M.J., and Wehbe, L. (2023). Better models of human high-level visual cortex emerge from natural language supervision with a large and diverse dataset. Nature Machine Intelligence 5, 1415–1426.

31. Du, C., Fu, K., Wen, B., Sun, Y., Peng, J., Wei, W., Gao, Y., Wang, S., Zhang, C., and Li, J. (2025). Human-like object concept representations emerge naturally in multimodal large language models. Nature Machine Intelligence, 1-16.

32. Jang, H., and Tong, F. (2024). Improved modeling of human vision by incorporating robustness to blur in convolutional neural networks. Nat Commun 15, 1989. 10.1038/s41467-024-45679-0.

33. Kubilius, J., Schrimpf, M., Nayebi, A., Bear, D., Yamins, D.L., and DiCarlo, J.J. (2018). Cornet: Modeling the neural mechanisms of core object recognition. BioRxiv, 408385.

34. Spoerer, C.J., McClure, P., and Kriegeskorte, N. (2017). Recurrent Convolutional Neural Networks: A Better Model of Biological Object Recognition. Frontiers in psychology 8, 1551. 10.3389/fpsyg.2017.01551.

35. Miller, G.A. (1995). WordNet: a lexical database for English. Communications of the ACM 38, 39–41.

36. Geirhos, R., Meding, K., and Wichmann, F.A. (2020). Beyond accuracy: quantifying trial-by-trial behaviour of CNNs and humans by measuring error consistency. Advances in neural information processing systems 33, 13890–13902.

37. Kubilius, J., Schrimpf, M., Kar, K., Rajalingham, R., Hong, H., Majaj, N., Issa, E., Bashivan, P., Prescott-Roy, J., and Schmidt, K. (2019). Brain-like object recognition with high-performing shallow recurrent ANNs. Advances in neural information processing systems 32.

38. Kar, K., Kubilius, J., Schmidt, K., Issa, E.B., and DiCarlo, J.J. (2019). Evidence that recurrent circuits are critical to the ventral stream’s execution of core object recognition behavior. Nature Neuroscience 22, 974–983. 10.1038/s41593-019-0392-5.

39. Tang, H., Schrimpf, M., Lotter, W., Moerman, C., Paredes, A., Ortega Caro, J., Hardesty, W., Cox, D., and Kreiman, G. (2018). Recurrent computations for visual pattern completion. Proceedings of the National Academy of Sciences 115, 8835–8840. doi:10.1073/pnas.1719397115.

40. Rajaei, K., Mohsenzadeh, Y., Ebrahimpour, R., and Khaligh-Razavi, S.M. (2019). Beyond core object recognition: Recurrent processes account for object recognition under occlusion. PLoS Comput Biol 15, e1007001. 10.1371/journal.pcbi.1007001.

41. Kar, K., and DiCarlo, J.J. (2021). Fast Recurrent Processing via Ventrolateral Prefrontal Cortex Is Needed by the Primate Ventral Stream for Robust Core Visual Object Recognition. Neuron 109, 164–176.e165. 10.1016/j.neuron.2020.09.035.

42. Dosovitskiy, A., Beyer, L., Kolesnikov, A., Weissenborn, D., Zhai, X., Unterthiner, T., Dehghani, M., Minderer, M., Heigold, G., and Gelly, S. (2020). An image is worth 16×16 words: Transformers for image recognition at scale. arXiv preprint arXiv:2010.11929.

43. Dong, X., Bao, J., Zhang, T., Chen, D., Gu, S., Zhang, W., Yuan, L., Chen, D., Wen, F., and Yu, N. (2022). Clip itself is a strong fine-tuner: Achieving 85.7% and 88.0% top-1 accuracy with vit-b and vit-l on imagenet. arXiv preprint arXiv:2212.06138.

44. Schrimpf, M., Kubilius, J., Hong, H., Majaj, N.J., Rajalingham, R., Issa, E.B., Kar, K., Bashivan, P., Prescott-Roy, J., and Geiger, F. (2018). Brain-score: Which artificial neural network for object recognition is most brain-like? BioRxiv, 407007.

45. Schrimpf, M., Kubilius, J., Lee, M.J., Ratan Murty, N.A., Ajemian, R., and DiCarlo, J.J. (2020). Integrative Benchmarking to Advance Neurally Mechanistic Models of Human Intelligence. Neuron 108, 413–423. 10.1016/j.neuron.2020.07.040.

46. DiCarlo, J.J., Zoccolan, D., and Rust, N.C. (2012). How does the brain solve visual object recognition? Neuron 73, 415–434. 10.1016/j.neuron.2012.01.010.

47. Hendrycks, D., and Dietterich, T. (2019). Benchmarking neural network robustness to common corruptions and perturbations. arXiv preprint arXiv:1903.12261.

48. Ahlert, J., Klein, T., Wichmann, F., and Geirhos, R. (2024). How aligned are different alignment metrics? arXiv preprint arXiv:2407.07530.

49. Freud, E., Culham, J.C., Plaut, D.C., and Behrmann, M. (2017). The large-scale organization of shape processing in the ventral and dorsal pathways. Elife 6. 10.7554/eLife.27576.

50. Bar, M., Kassam, K.S., Ghuman, A.S., Boshyan, J., Schmid, A.M., Dale, A.M., Hämäläinen, M.S., Marinkovic, K., Schacter, D.L., Rosen, B.R., and Halgren, E. (2006). Top-down facilitation of visual recognition. Proceedings of the National Academy of Sciences 103, 449–454. 10.1073/pnas.0507062103.

51. Fyall, A.M., El-Shamayleh, Y., Choi, H., Shea-Brown, E., and Pasupathy, A. (2017). Dynamic representation of partially occluded objects in primate prefrontal and visual cortex. Elife 6. 10.7554/eLife.25784.

52. Wang, M., Arteaga, D., and He, B.J. (2013). Brain mechanisms for simple perception and bistable perception. Proc Natl Acad Sci U S A 110, E3340–E3349. 10.1073/pnas.1221945110.

53. Bracci, S., and Op de Beeck, H. (2016). Dissociations and Associations between Shape and Category Representations in the Two Visual Pathways. J Neurosci 36, 432–444. 10.1523/JNEUROSCI.2314-15.2016.

54. Levinson, M., Podvalny, E., Baete, S.H., and He, B.J. (2021). Cortical and subcortical signatures of conscious object recognition. Nature Communications 12, 2930. 10.1038/s41467-021-23266-x.

55. Ayzenberg, V., Simmons, C., and Behrmann, M. (2023). Temporal asymmetries and interactions between dorsal and ventral visual pathways during object recognition. Cerebral cortex communications 4, tgad003. 10.1093/texcom/tgad003.

56. Konen, C.S., and Kastner, S. (2008). Two hierarchically organized neural systems for object information in human visual cortex. Nat Neurosci 11, 224–231. 10.1038/nn2036.

57. Doerig, A., Kietzmann, T.C., Allen, E., Wu, Y., Naselaris, T., Kay, K., and Charest, I. (2025). High-level visual representations in the human brain are aligned with large language models. Nature Machine Intelligence, 1–15.

58. Potter, M.C., Wyble, B., Hagmann, C.E., and McCourt, E.S. (2014). Detecting meaning in RSVP at 13 ms per picture. Atten Percept Psychophys 76, 270–279. 10.3758/s13414-013-0605-z.

59. Rosch, E., Mervis, C.B., Gray, W.D., Johnson, D.M., and Boyes-Braem, P. (1976). Basic objects in natural categories. Cognitive psychology 8, 382–439.

60. Mehrer, J., Spoerer, C.J., Jones, E.C., Kriegeskorte, N., and Kietzmann, T.C. (2021). An ecologically motivated image dataset for deep learning yields better models of human vision. Proceedings of the National Academy of Sciences 118, e2011417118. 10.1073/pnas.2011417118.

61. Muttenthaler, L., Greff, K., Born, F., Spitzer, B., Kornblith, S., Mozer, M.C., Müller, K.-R., Unterthiner, T., and Lampinen, A.K. (2025). Aligning machine and human visual representations across abstraction levels. Nature 647, 349–355. 10.1038/s41586-025-09631-6.

62. Schmidt, F., Hebart, M.N., Schmid, A.C., and Fleming, R.W. (2025). Core dimensions of human material perception. Proceedings of the National Academy of Sciences 122, e2417202122.

63. Henderson, M.M., Tarr, M.J., and Wehbe, L. (2023). A texture statistics encoding model reveals hierarchical feature selectivity across human visual cortex. Journal of Neuroscience 43, 4144–4161.

64. Casile, A., Cordier, A., Kim, J.G., Cometa, A., Madsen, J.R., Stone, S., Ben-Yosef, G., Ullman, S., Anderson, W., and Kreiman, G. (2025). Neural correlates of minimal recognizable configurations in the human brain. Cell reports 44.

65. Ayzenberg, V., and Behrmann, M. (2022). Does the brain’s ventral visual pathway compute object shape? Trends Cogn Sci 26, 1119–1132. 10.1016/j.tics.2022.09.019.

66. Cherian, T., Jacob, G., and Arun, S. (2025). Do monkeys see the way we do? Qualitative similarities and differences between monkey and human perception. bioRxiv, 2025.2001. 2030.635614.

67. Russakovsky, O., Deng, J., Su, H., Krause, J., Satheesh, S., Ma, S., Huang, Z., Karpathy, A., Khosla, A., and Bernstein, M. (2015). Imagenet large scale visual recognition challenge. International journal of computer vision 115, 211–252.

68. Kubilius, J., Bracci, S., and Op de Beeck, H.P. (2016). Deep Neural Networks as a Computational Model for Human Shape Sensitivity. PLoS Comput Biol 12, e1004896. 10.1371/journal.pcbi.1004896.

69. Gatys, L., Ecker, A.S., and Bethge, M. (2015). Texture synthesis using convolutional neural networks. Advances in neural information processing systems 28.

70. Gatys, L.A., Ecker, A.S., and Bethge, M. (2016). Image style transfer using convolutional neural networks. pp. 2414–2423.

71. Chan, C., Durand, F., and Isola, P. (2022). Learning to generate line drawings that convey geometry and semantics. pp. 7915–7925.

72. Singer, J.J.D., Cichy, R.M., and Hebart, M.N. (2023). The Spatiotemporal Neural Dynamics of Object Recognition for Natural Images and Line Drawings. J Neurosci 43, 484–500. 10.1523/JNEUROSCI.1546-22.2022.

73. Gonzalez-Garcia, C., Flounders, M.W., Chang, R., Baria, A.T., and He, B.J. (2018). Content-specific activity in frontoparietal and default-mode networks during prior-guided visual perception. eLife 7. 10.7554/eLife.36068.

74. Paszke, A. (2019). Pytorch: An imperative style, high-performance deep learning library. arXiv preprint arXiv:1912.01703.

75. Liu, Z., Mao, H., Wu, C.-Y., Feichtenhofer, C., Darrell, T., and Xie, S. (2022). A convnet for the 2020s. pp. 11976–11986.

76. Schuhmann, C., Beaumont, R., Vencu, R., Gordon, C., Wightman, R., Cherti, M., Coombes, T., Katta, A., Mullis, C., and Wortsman, M. (2022). Laion-5B: An open large-scale dataset for training next generation image-text models. Advances in neural information processing systems 35, 25278–25294.

77. Ilharco, G., Wortsman, M., Wightman, R., Gordon, C., Carlini, N., et al. (2025). OpenCLIP. Zenodo. 10.5281/zenodo.5143772.

78. Wightman, R., Raw, N., Soare, A., Arora, A., Ha, C., and Christoph Reich (2023). pytorch-image-models. Zenodo. 10.5281/zenodo.4414861.

79. Li, A.C., Prabhudesai, M., Duggal, S., Brown, E., and Pathak, D. (2023). Your diffusion model is secretly a zero-shot classifier. pp. 2206–2217.

80. Rajalingham, R., Issa, E.B., Bashivan, P., Kar, K., Schmidt, K., and DiCarlo, J.J. (2018). Large-Scale, High-Resolution Comparison of the Core Visual Object Recognition Behavior of Humans, Monkeys, and State-of-the-Art Deep Artificial Neural Networks. The Journal of Neuroscience 38, 7255–7269. 10.1523/jneurosci.0388-18.2018.

81. Mahner, F.P., Muttenthaler, L., Guclu, U., and Hebart, M.N. (2025). Dimensions underlying the representational alignment of deep neural networks with humans. Nat Mach Intell 7, 848–859. 10.1038/s42256-025-01041-7.

