## Supplemental figures for "Systematic image perturbations reveal persistent gaps between human and machine vision"

### SUPPLEMENTARY INFORMATION

**A**

| Animate | | Inanimate | |
|---|---|---|---|
| **Mammal** | **Non-mammal** | **Man-made** | **Natural** |
| Bear | Bird | Airplane | Pineapple |
| Leporid | Snake | Car | Banana |
| Cat | Frog | Chair | Flower |
| Dog | Crab | Shoe | Broccoli |
| Elephant | Butterfly | Clock | Cabbage |
| Monkey | Fish | Microwave | Strawberry |
| Tiger | Crocodilian | Gun | Bell pepper |
| Camel | Turtle | Guitar | Mushroom |
| Swine | Lobster | Mug | Lemon |
| Horse | Bee | Dryer | Acorn |
| Hippopotamus | Spider | Helmet | Corn |
| Squirrel | Snail | Hammer | Apple |

**C**

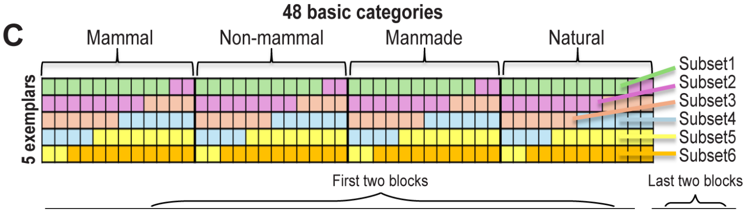


| | Subset1 | Subset2 | Subset3 | Subset4 | Subset5 | Subset6 | | Full set |
|---|---|---|---|---|---|---|---|---|
| Subject1 | Fil | Lin | SilTex | Sil | Tex | LinFll | | Ori |
| Subject2 | LinFll | Fil | Lin | SilTex | Sil | Tex | | Ori |
| Subject3 | Tex | LinFll | Fil | Lin | SilTex | Sil | | Ori |
| Subject4 | Sil | Tex | LinFll | Fil | Lin | SilTex | + | Ori |
| Subject5 | SilTex | Sil | Tex | LinFll | Fil | Lin | | Ori |
| Subject6 | Lin | SilTex | Sil | Tex | LinFll | Fil | | Ori |
| ⋮ | | | ⋮ | | | | | ⋮ |

Pooled subject1; Pooled subject2; ⋮

*The image subsets were regenerated for every group of 6 subjects.

**B**

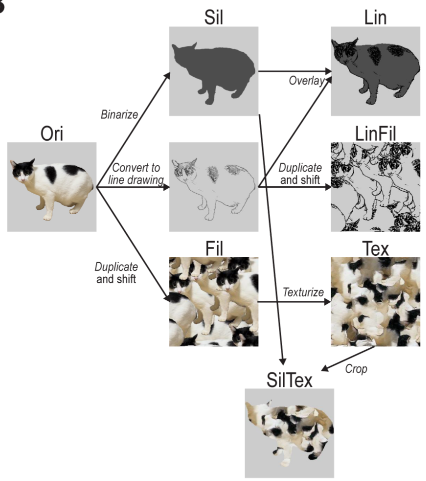


**D**

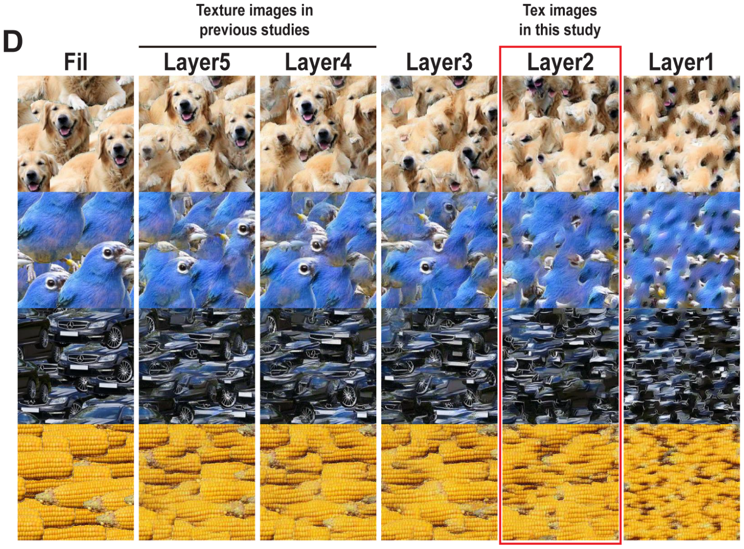


**Fig. S1. Image set construction and detailed experimental design.** (**A**) A list of 48 basic categories included. We defined these 48 categories by selecting 12 categories from each of four coarse groups (i.e., mammals, non-mammal animals, man-made objects, and natural objects) to create a broad stimulus set. (**B**) Workflow for generating each image condition in Experiment 1 (see Methods for details). (**C**) Each cell in the top panel represents an exemplar image and is color-coded to show the subset to which it belongs. The 240 exemplars were partitioned into six subsets of 40 images each, with each subset containing one exemplar from 40 basic categories—10 drawn from each of the four coarse groups. Each subject viewed one of the six subsets under a given manipulation, with assignments of subset-to-manipulation counterbalanced across the six subjects within each group. As such, collectively across the six subjects in the same group, every exemplar image was presented in every condition and each subject only saw a given exemplar image (e.g., blue jay picture) in a single manipulation condition. There were 10 groups of six subjects, resulting in 10 pooled subjects using data from 60 individual subjects. (**D**) Results of texturization across different layer sets. The leftmost column shows example Fil images. The five columns to the right show images generated by matching the Gram matrix of each Fil image using layer sets that include all layers up to pooling layers 1 through 5, respectively.

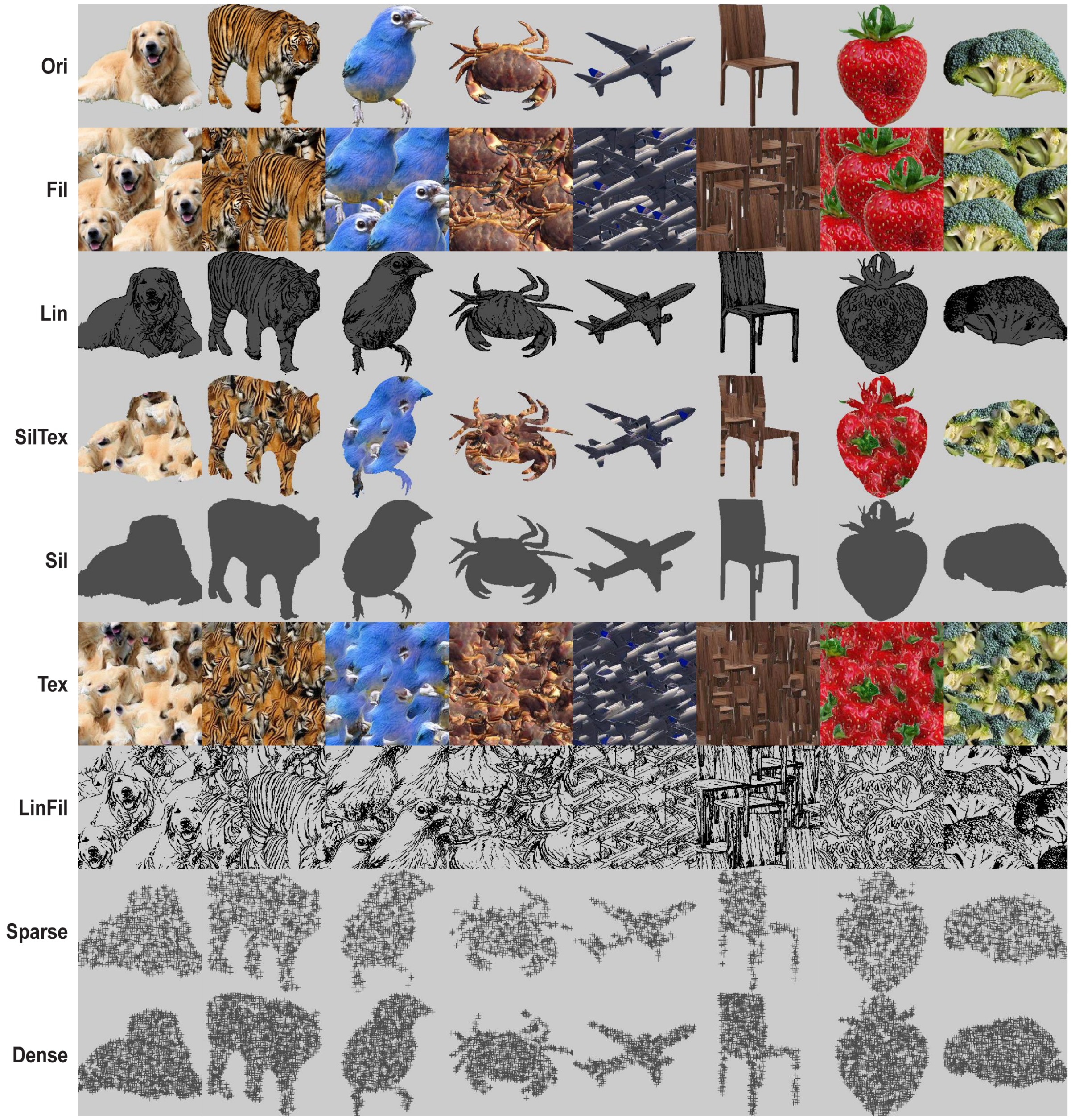


**Fig. S2. Additional example images for each image condition.** Due to copyright restrictions, the exemplars are replaced with publicly available, copyright-free images that closely resemble the actual images used in the experiment.

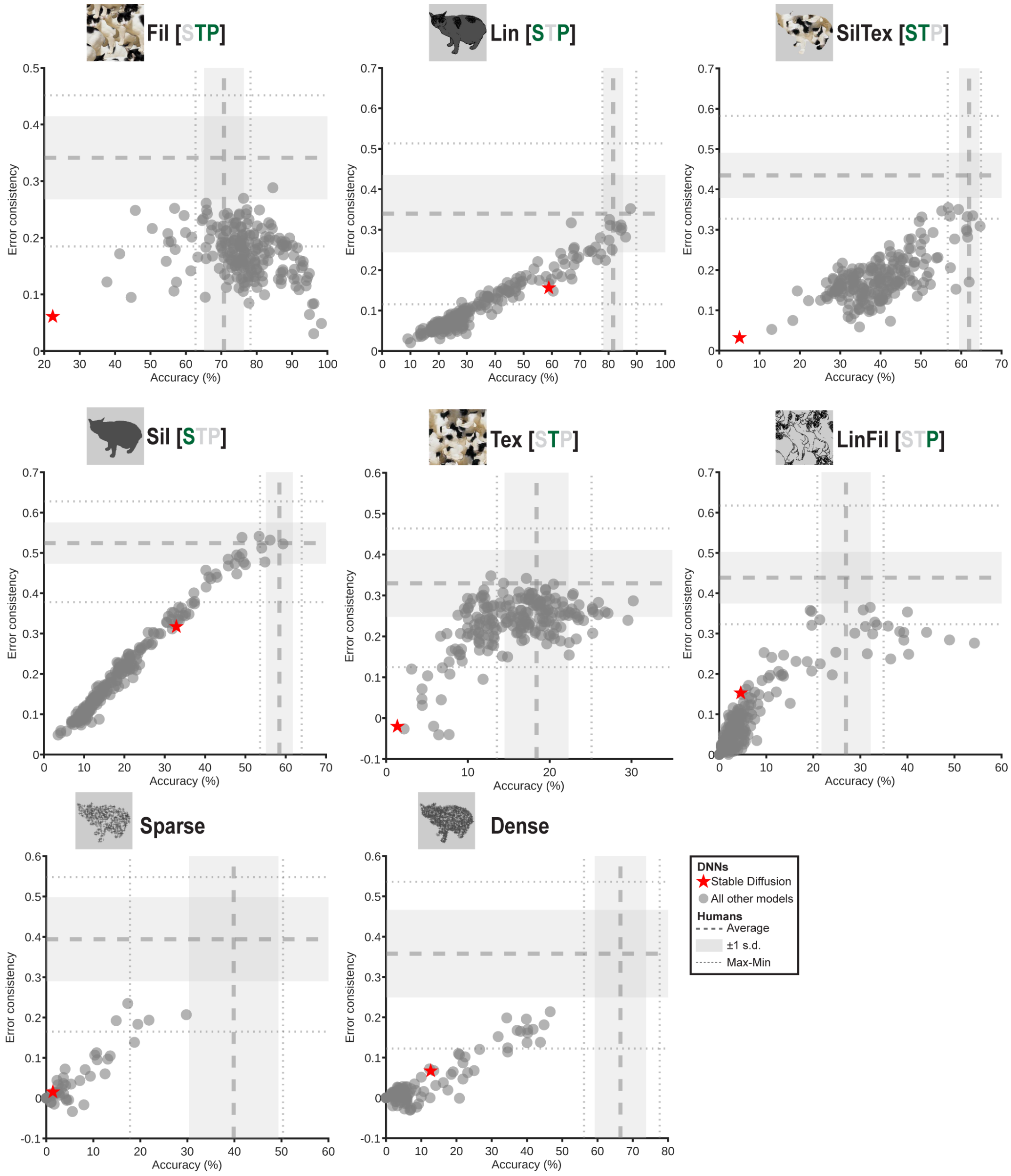


**Fig. S3. Performance of the stable diffusion model.** Red stars denote recognition accuracy and error consistency with humans for the Stable Diffusion classifier across the six conditions in Experiment 1 and the two conditions in Experiment 2. Gray dots indicate all other 209 DNNs, reproduced from Figs. 3 and 4.

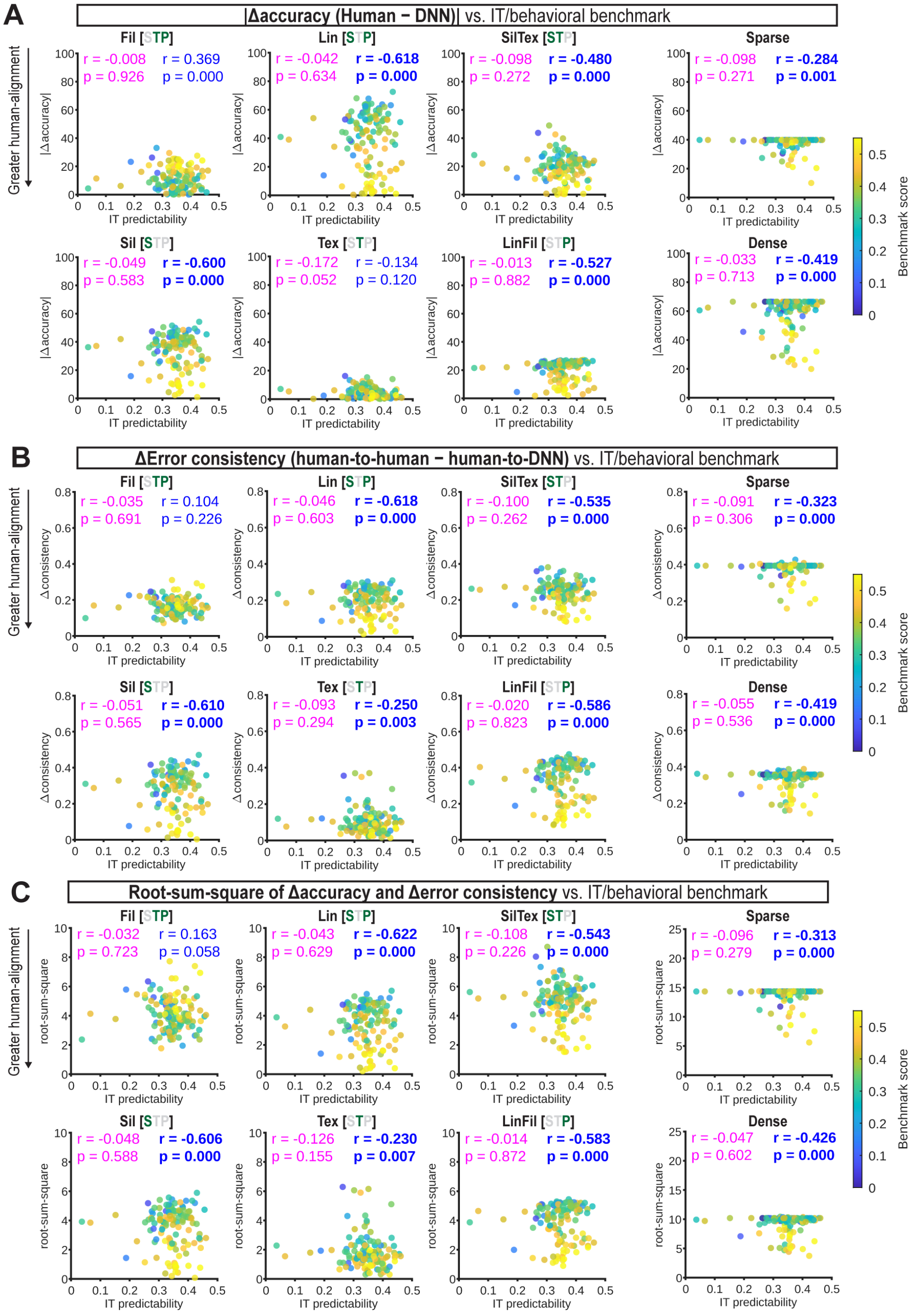


**Fig. S4. Correlation between human alignment in our image set and neural and behavioral benchmarks in the Brain-Score.** This analysis includes 128 of 209 DNN models for which IT predictability scores were available in the Brain-Score database. Each dot represents one DNN, with dot color denoting the mean Brain-Score behavioral benchmark score for that DNN, as indicated by the color bar on the right. Abscissa values in all plots represent IT-cortex neural predictability scores from the Brain-Score database. The ordinate values represent (**A**) absolute difference in recognition accuracy from humans, (**B**) difference between human-to-human and human-to-DNN error consistency, and (**C**) root-sum-of-square distance ($D_m$) from human average in the two-dimensional space defined by accuracy and error consistency after z-score normalization (see Methods). Magenta annotations indicate correlation between IT-cortex neural predictability scores (abscissa values) and the ordinate values. Blue annotations indicate correlations between the mean Brain-Score behavioral benchmark score (as indicated by dot color) and the ordinate values. Bold font indicates significant ($p < 0.05$, uncorrected) *negative* correlations, which indicate that Brain-Score alignment positively correlates with human alignment as assessed by our image set.

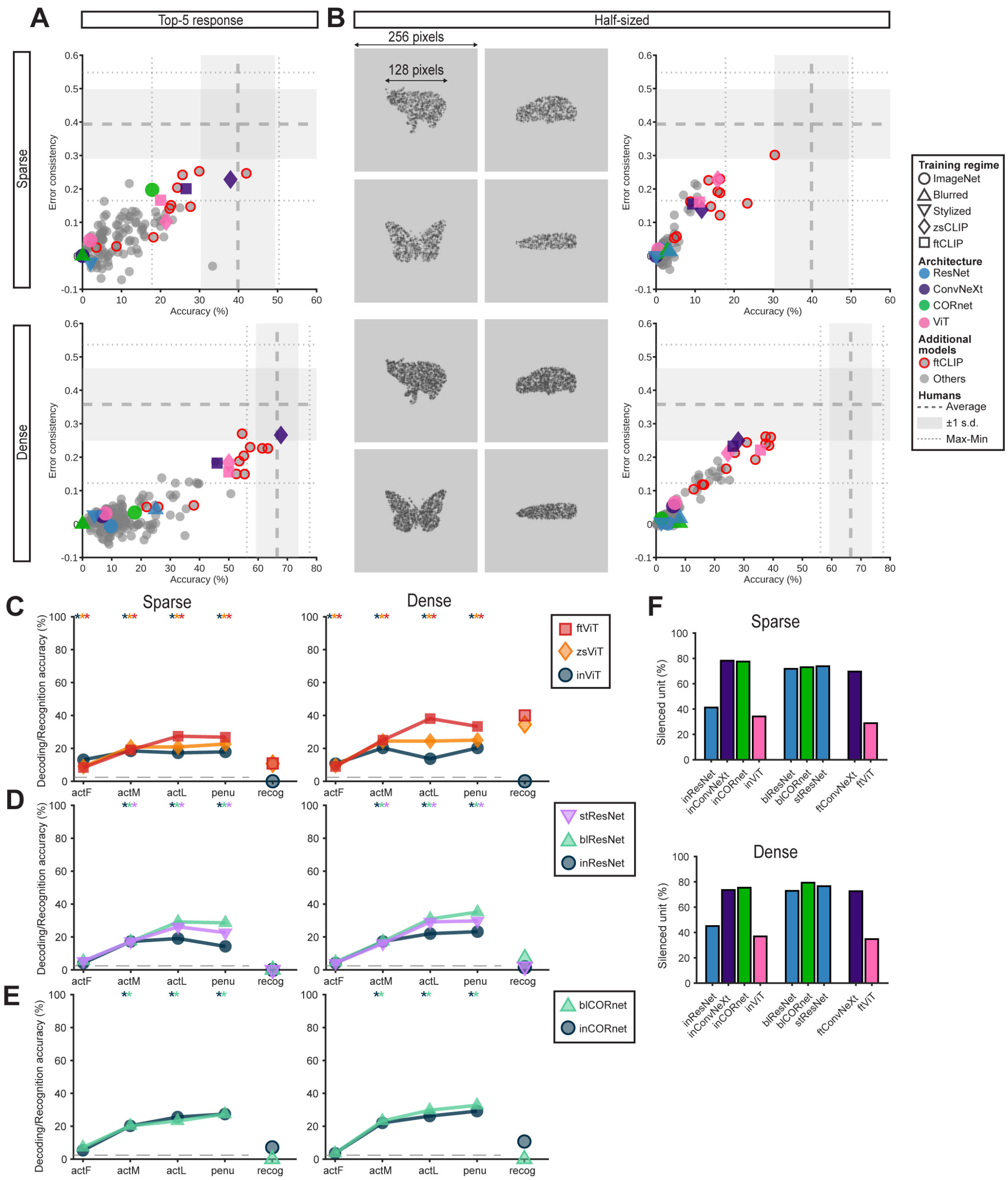


**Fig. S5. Additional analyses for Sparse and Dense image recognition, decoding accuracy for ViT-, ResNet50-, and CORnet-S-based models in Experiment 2, and percentage of locally tuned units.** (**A**) The results corresponding to the bottom of Fig. 4, but using DNNs' top-5 predictions. (**B**) Example half-sized **Sparse** and **Dense** images (left) and corresponding results (right). As checkpoints for six models became unavailable during the study, this analysis includes 203 of the 209 DNNs. (**C–E**) Decoding accuracy of the ViT- (**C**), ResNet50- (**D**), and CORnet-S-based (**E**) models at each of the four representative layers in the **Sparse** (left) and **Dense** (right) conditions. Dashed lines indicate chance level (1/41). Stars above at the top indicate significant differences from chance (label permutation test; $p < 0.05$; Bonferroni-corrected over layers; see Methods for details). The rightmost "recog" column shows recognition accuracy reproduced from Fig. 4 for comparison. (**F**) Percentage of locally tuned units in the **Sparse** (top) or **Dense** (bottom) conditions.

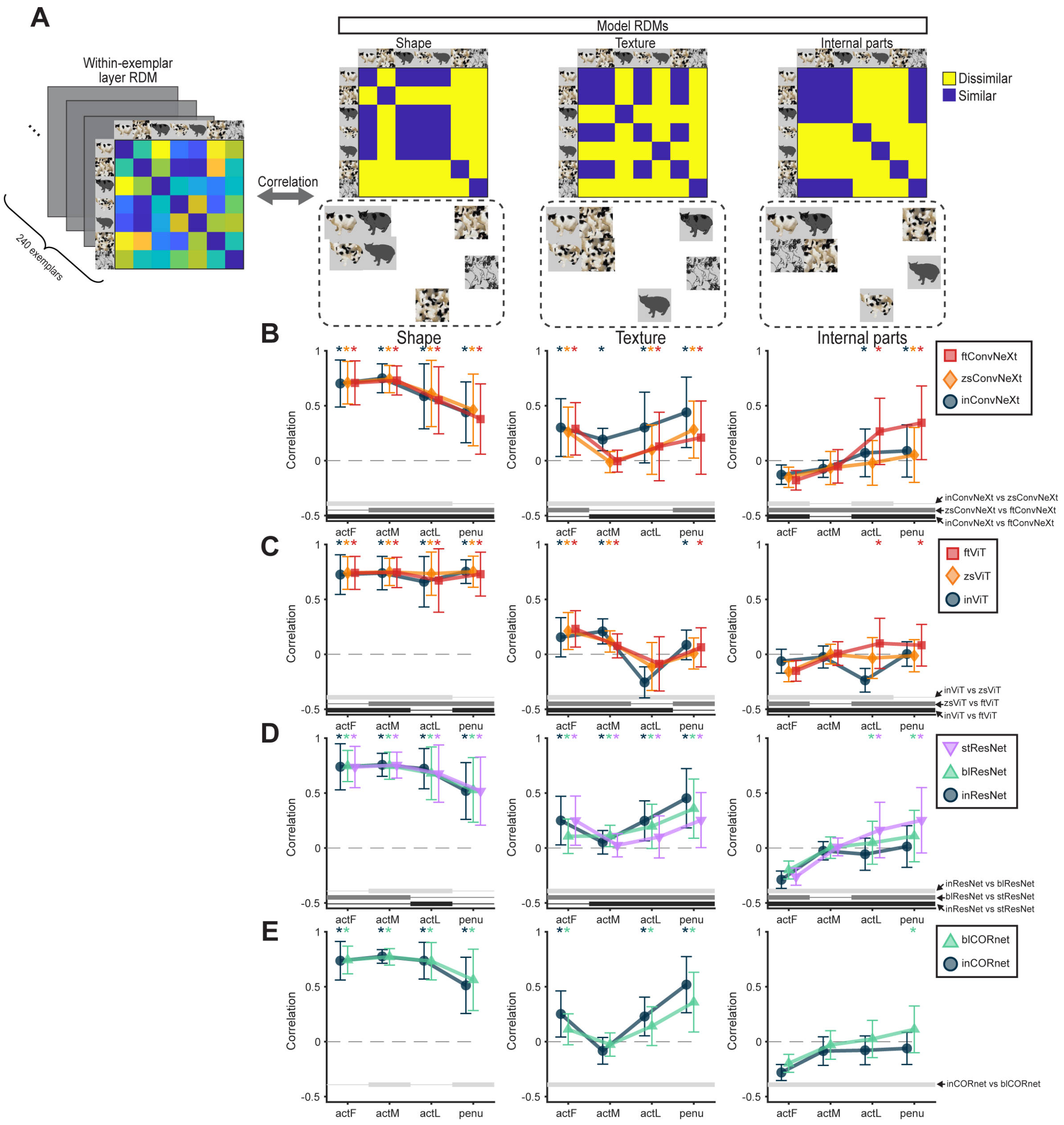


**Fig. S6. Layer-wise dynamics of shape, texture, and internal-parts representations.** (**A**) Exemplar-wise RSA schematic. For each exemplar image, we first computed a representational dissimilarity matrix (RDM) by measuring the correlational distance among activation patterns evoked by the seven image conditions (left). Each layer's RDM was correlated with three model RDMs hypothesizing representation of shape, texture, or internal parts (right). Below each RDM, a two-dimensional embedding illustrates the approximate relative distances among the seven conditions. (**B–E**) Spearman's rank correlation between each layer's RDM and the shape (left), texture (middle), and internal-parts (right) RDMs, averaged across exemplars in ConvNext- (**B**) ViT- (**C**), ResNet50- (**D**), and CORnet-S-based (**E**) models. Error bars denote standard deviation across exemplar images. Bottom thick horizontal lines indicate two-sided paired t-test between model pairs (**b**: top: inConvNeXt vs. zsConvNeXt; middle: zsConvNeXt vs. ftConvNeXt; bottom: inConvNeXt vs. ftConvNeXt; **C**: top: inViT vs. zsViT; middle: zsViT vs. ftViT; bottom: inViT vs. ftViT; **D**: top: inResNet vs. blResNet; middle: blResNet vs. stResNet; bottom: inResNet vs. stResNet; **E**: inCORnet vs. blCORnet; N=240; $p < 0.05$; Bonferroni-corrected over layers). Stars at the top denote significant one-sided t-tests against baseline zero ($N = 240$; $p < 0.05$; Bonferroni-corrected over layers).

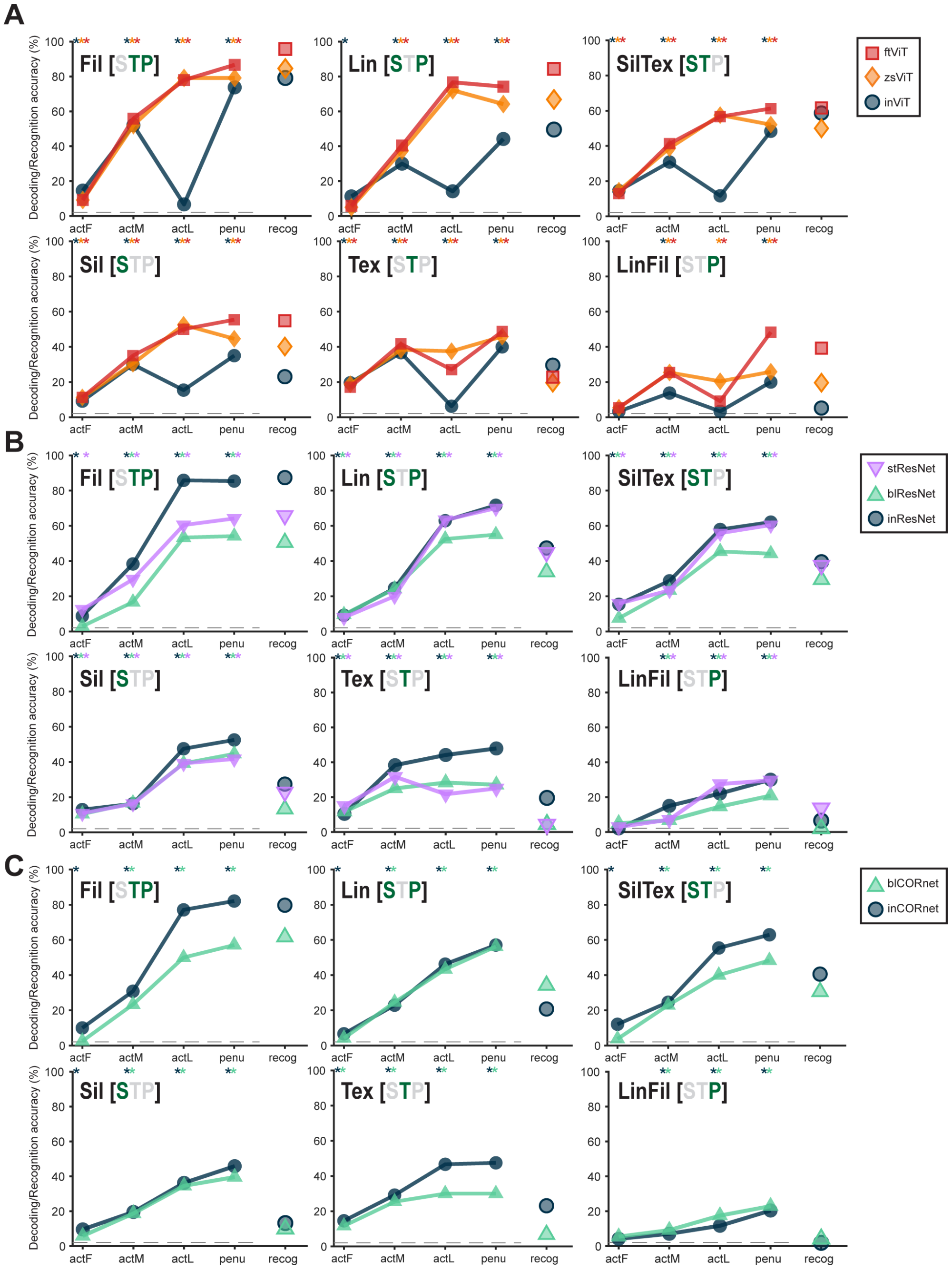


**Fig. S7. Decoding accuracy for ViT-, ResNet50-, and CORnet-S-based models in Experiment 1.** (**A–C**) Decoding accuracy of the ViT- (**A**), ResNet50- (**B**), and CORnet-S-based (**C**) models at each of the four representative layers across conditions. Dashed line indicates chance level (1/48). Stars at the top indicate significant differences from chance (label permutation test; $p < 0.05$; Bonferroni-corrected over layers; see Methods for details; exact statistics in the supplementary Excel file). The rightmost “recog” column shows recognition accuracy reproduced from Fig. 2 for comparison.

**Table S1. List of 198 DNNs included in large-scale analyses.** This table provides the "model identifier" for each of the 198 DNNs, as defined in the Brain-Score database (https://github.com/brain-score/vision).

| | | |
|---|---|---|
| AT_efficientnet-b2 | eBarlow_augself_linear_1 | mobilenet_v2_1_0_224 |
| AdvProp_efficientnet-b4 | eBarlow_augself_mlp_1 | mobilenet_v2_1_3_224 |
| AdvProp_efficientnet-b6 | eBarlow_lmda_0001_1 | mobilenet_v2_1_4_224 |
| AdvProp_efficientnet-b7 | eBarlow_lmda_001_1 | nasnet_large |
| AdvProp_efficientnet-b8 | eBarlow_lmda_001_2 | nasnet_mobile |
| AlexNet_SIN | eBarlow_lmda_001_3 | pnasnet_large |
| BiT-S-R101x1 | eBarlow_lmda_01_1 | r101_eBarlow_Vanilla_1 |
| BiT-S-R101x3 | eBarlow_lmda_01_2 | r101_eBarlow_lmda_01_1 |
| BiT-S-R152x2 | eBarlow_lmda_01 | r101_eBarlow_lmda_02_1 |
| BiT-S-R152x4 | eBarlow_lmda_02_1000ep | r34_eMMCR_Mom_Vanilla_1 |
| BiT-S-R50x1 | eBarlow_lmda_02_1 | r34_eMMCR_Mom_lmda_01_1 |
| BiT-S-R50x3 | eBarlow_lmda_02_200_full | r34_eMMCR_Mom_lmda_02_1 |
| CORnet-S | eBarlow_lmda_03_1 | r50_tvpt |
| ReAlnet01 | eBarlow_lmda_04_1 | regnet |
| ReAlnet02 | eBarlow_lmda_05_1 | resnet101_imagenet_full |
| ReAlnet03 | eMMCR_Mom_Vanilla_1 | resnet152_imagenet_full |
| ReAlnet04 | eMMCR_Mom_Vanilla_2 | resnet18_imagenet_full |
| ReAlnet05 | eMMCR_Mom_lmda_0001_1 | resnet34_imagenet_full |
| ReAlnet06 | eMMCR_Mom_lmda_001_1 | resnet50-SIN_IN_IN |
| ReAlnet07 | eMMCR_Mom_lmda_01_1 | resnet50-SIN_IN |
| ReAlnet08 | eMMCR_Mom_lmda_01_2 | resnet50-SIN |
| ReAlnet09 | eMMCR_Mom_lmda_02_1 | resnet50-VITO-8deg-cc |
| ReAlnet10 | eMMCR_Mom_lmda_03_1 | resnet50-moclr8deg |
| Res2Net50_26w_4s | eMMCR_Mom_lmda_04_1 | resnet50_eMMCR_VanillaV2 |
| VOneCORnet-S | eMMCR_Mom_lmda_05_1 | resnet50_eMMCR_eqp10_lm1V2 |
| ViT_L_32_imagenet1k | eMMCR_VanillaV2 | resnet50_finetune_cutmix_AVGe2e3_robust_linf8255_e0_247x234 |
| alexnet_7be5be79 | eMMCR_Vanilla_1 | resnet50_imagenet_100_seed-0 |
| antialias-resnet152 | eMMCR_Vanilla_2 | resnet50_imagenet_full |
| antialiased-r50 | eMMCR_lmda_01V2 | resnet50_robust_l2_eps1 |
| antialiased-rnext101_32x8d | eMMCR_lmda_01_1 | resnet_101_v1 |
| convnext_tiny:in12k_ft_in1k | eMMCR_lmda_01_2 | resnet_101_v2 |
| convnext_xlarge:fb_in22k_ft_in1k | eSimCLR_Vanilla_1 | resnet_152_v1 |
| convnext_xxlarge:clip_laion2b_soup_ft_in1k | eSimCLR_Vanilla_2 | resnet_152_v2 |
| vit_base_patch16_clip_224:openai_ft_in12k_in1k | eSimCLR_lmda_0001_1 | resnet_50_2 |
| vit_huge_patch14_clip_224:laion2b_ft_in12k_in1k | eSimCLR_lmda_001_1 | resnet_50_v2 |
| vit_large_patch14_clip_224:laion2b_ft_in12k_in1k | eSimCLR_lmda_01_1 | resnext101_32x16d_wsl |
| vit_large_patch14_clip_224:openai_ft_in12k_in1k | eSimCLR_lmda_01_2 | resnext101_32x32d_wsl |
| vit_large_patch14_clip_336:openai_ft_in12k_in1k | eSimCLR_lmda_02_1_1 | resnext101_32x48d_wsl |
| bp_resnet50_julios | eSimCLR_lmda_03_1 | resnext101_32x8d_wsl |
| convnext_base:clip_laiona_augreg_ft_in1k_384 | eSimCLR_lmda_04_1_1 | sBarlow_lmda_01 |
| convnext_base_imagenet_full_seed-0 | eSimCLR_lmda_05_1 | sBarlow_lmda_0 |
| convnext_femto_ols:d1_in1k | efficientnet-b7 | sBarlow_lmda_1 |
| convnext_large:fb_in22k_ft_in1k | efficientnet_b0 | sBarlow_lmda_2 |
| convnext_large_imagenet_full_seed-0 | efficientnet_b0_imagenet_full | sBarlow_lmda_8 |
| convnext_small_imagenet_100_seed-0 | efficientnet_b1_imagenet_full | shufflenet_v2_x1_0 |
| convnext_small_imagenet_full_seed-0 | efficientnet_b2_imagenet_full | swin_small_patch4_window7_224:ms_in22k_ft_in1k |
| convnext_tiny_imagenet_full_seed-0 | effnetb1_272x240 | tv_efficientnet-b1 |
| convnext_tiny_sup | effnetb1_cutmix_augmix_sam_e1_5avg_424x377 | vgg_16 |
| custom_model_cv_18_dagger_408 | effnetb1_cutmixpatch_SAM_robust32_avge6e8e9e10_manylayers_324x288 | vgg_19 |
| cv_18_dagger_408_pretrained | effnetb1_cutmixpatch_augmix_robust32_avge4e7_manylayers_324x288 | vit_base_patch16_clip_224:openai_ft_in1k |
| cvt_cvt-13-224-in1k_4_LucyV4 | evresnet_50_4 | vit_huge_patch14_clip_336:laion2b_ft_in12k_in1k |
| cvt_cvt-13-384-in1k_4_LucyV4 | fixres_resnext101_32x48d_wsl | vit_large_patch14_clip_224:laion2b_ft_in1k |
| cvt_cvt-13-384-in22k_finetuned-in1k_4_LucyV4 | focalnet_tiny_lrf_in1k | vit_large_patch14_clip_224:openai_ft_in1k |
| cvt_cvt-21-224-in1k_4_LucyV4 | grcnn_109 | vit_large_patch14_clip_336:laion2b_ft_in1k |
| cvt_cvt-21-384-in1k_4_LucyV4 | grcnn | vit_relpos_base_patch16_clsgap_224:sw_in1k |
| cvt_cvt-21-384-in22k_finetuned-in1k_4_LucyV4 | imagenet_l2_3_0 | vit_relpos_base_patch32_plus_rpn_256:sw_in1k |
| cvt_cvt-w24-384-in22k_finetuned-in1k_4 | inception_v1 | vit_tiny_r_s16_p8_384:augreg_in21k_ft_in1k |
| dbp_resnet50_julios | inception_v3 | vonegrcnn_47e |
| deit_base_imagenet_full_seed-0 | inception_v4 | vonegrcnn_52e_full |
| deit_large_imagenet_full_seed-0 | mobilenet_v2_0_5_192 | vonegrcnn_62e_nobn |
| deit_small_imagenet_full_seed-0 | mobilenet_v2_0_5_224 | voneresnet-50-non_stochastic |
| densenet-121 | mobilenet_v2_0_75_160 | voneresnet-50-robust |
| densenet-169 | mobilenet_v2_0_75_192 | voneresnet-50 |
| densenet-201 | mobilenet_v2_0_75_224 | voneresnet_50_1 |
| eBarlow_Vanilla_1 | mobilenet_v2_1_0_128 | xception |
| eBarlow_Vanilla_2 | mobilenet_v2_1_0_160 | |
| eBarlow_Vanilla | mobilenet_v2_1_0_192 | |

**Table S2. Top 10 DNNs that were close to human behavior in each condition.** For each condition, after z-score normalization we computed the Euclidean distance $D_m$ between each model and the human average in the two-dimensional space defined by normalized accuracy and normalized error consistency (see "Correlation between IT predictability and behavioral similarities to humans" in Methods). The top 10 DNNs with the smallest $D_m$ in each condition are listed; pink shading indicates fine-tuned CLIP models, and green shading indicates zero-shot CLIP models.

| Fil | Lin | SilTex |
|---|---|---|
| cvt_cvt-21-384-in22k_finetuned-in1k_4_LucyV4 | cvt_cvt-w24-384-in22k_finetuned-in1k_4 | vit_large_patch14_clip_224:laion2b_ft_in1k |
| clip_vit-l-laion_image | vit_huge_patch14_clip_224:laion2b_ft_in12k_in1k | vit_large_patch14_clip_224:openai_ft_in12k_in1k |
| mobilenet_v2_0_75_192 | convnext_large_mlp:clip_laion2b_augreg_ft_in1k_384 | vit_large_patch14_clip_224:openai_ft_in1k |
| resnet50-sin | vit_large_patch14_clip_224:laion2b_ft_in12k_in1k | vit_huge_patch14_clip_224:laion2b_ft_in12k_in1k |
| ViT_L_32_imagenet1k | vit_huge_patch14_clip_336:laion2b_ft_in12k_in1k | vit_large_patch14_clip_336:openai_ft_in12k_in1k |
| eBarlow_lmda_05_1 | vit_large_patch14_clip_224:laion2b_ft_in1k | convnext_large_mlp:clip_laion2b_augreg_ft_in1k_384 |
| ReAlnet09 | vit_large_patch14_clip_224:openai_ft_in12k_in1k | clip_convnextL_image |
| xception | clip_vit-l-laion_image | vit_huge_patch14_clip_336:laion2b_ft_in12k_in1k |
| resnet50_robust_l2_eps1 | convnext_xxlarge:clip_laion2b_soup_ft_in1k | vit_large_patch14_clip_336:laion2b_ft_in1k |
| cvt_cvt-21-224-in1k_4_LucyV4 | resnext101_32x32d_wsl | resnext101_32x8d_wsl |

| Sil | Tex | LinFil |
|---|---|---|
| convnext_xxlarge:clip_laion2b_soup_ft_in1k | efficientnet_b0_imagenet_full | resnext101_32x48d_wsl |
| vit_huge_patch14_clip_224:laion2b_ft_in12k_in1k | convnext_xxlarge:clip_laion2b_soup_ft_in1k | clip_convnextL_image |
| vit_huge_patch14_clip_336:laion2b_ft_in12k_in1k | convnext_base:clip_laiona_augreg_ft_in1k_384 | vit_base_patch16_clip_224:openai_ft_in1k |
| vit_large_patch14_clip_224:openai_ft_in12k_in1k | eBarlow_lmda_01_1 | clip_vit-l-laion_image |
| vit_large_patch14_clip_224:laion2b_ft_in12k_in1k | clip_convnextL_image | cvt_cvt-w24-384-in22k_finetuned-in1k_4 |
| convnext_large_mlp:clip_laion2b_augreg_ft_in1k_384 | mobilenet_v2_1_0_128 | resnext101_32x32d_wsl |
| resnext101_32x32d_wsl | resnext101_32x32d_wsl | vit_large_patch14_clip_336:laion2b_ft_in1k |
| convnext_xlarge:fb_in22k_ft_in1k | deit_base_imagenet_full_seed-0 | vit_large_patch14_clip_224:laion2b_ft_in1k |
| vit_large_patch14_clip_224:laion2b_ft_in1k | Res2Net50_26w_4s | vit_base_patch16_clip_224:openai_ft_in12k_in1k |
| convnext_large:fb_in22k_ft_in1k | clip_vit-l-laion_image | resnext101_32x16d_wsl |

| Sparse | Dense |
|---|---|
| convnext_xxlarge:clip_laion2b_soup_ft_in1k | convnext_xxlarge:clip_laion2b_soup_ft_in1k |
| vit_huge_patch14_clip_224:laion2b_ft_in12k_in1k | vit_large_patch14_clip_336:openai_ft_in12k_in1k |
| clip_convnextL_image | clip_convnextL_image |
| convnext_large_mlp:clip_laion2b_augreg_ft_in1k_384 | AdvProp_efficientnet-b7 |
| vit_huge_patch14_clip_336:laion2b_ft_in12k_in1k | vit_huge_patch14_clip_224:laion2b_ft_in12k_in1k |
| vit_large_patch14_clip_224:laion2b_ft_in1k | vit_large_patch14_clip_224:openai_ft_in1k |
| vit_large_patch14_clip_336:openai_ft_in12k_in1k | vit_large_patch14_clip_224:laion2b_ft_in1k |
| vit_large_patch14_clip_224:openai_ft_in1k | vit_large_patch14_clip_336:laion2b_ft_in1k |
| clip_vit-l-laion_image | vit_huge_patch14_clip_336:laion2b_ft_in12k_in1k |
| vit_large_patch14_clip_336:laion2b_ft_in1k | vit_large_patch14_clip_224:openai_ft_in12k_in1k |

**Table S3. Labels for four representative hidden layers in each model.**

| | ActF | ActM | ActL | penu |
|---|---|---|---|---|
| **inResNet** | relu | layer3.1.relu | layer4.2.relu | avgpool |
| **inConvNeXt** | features.1.0.block.4 | features.5.2.block.4 | features.7.2.block.4 | classifier.0 |
| **inCORnet** | V1.nonlin1 | V4.nonlin1 | IT.nonlin3 | decoder.avgpool |
| **inViT** | encoder.layers.encoder_layer_0.mlp.1 | encoder.layers.encoder_layer_12.mlp.1 | encoder.layers.encoder_layer_23.mlp.1 | encoder.ln |
| **blResNet-blur** | relu | layer3.1.relu | layer4.2.relu | avgpool |
| **blCORnet** | V1.nonlin1 | V4.nonlin1 | IT.nonlin3 | decoder.avgpool |
| **stResNet** | relu | layer3.1.relu | layer4.2.relu | avgpool |
| **zsConvNeXt** | visual.trunk.stages.0.blocks.0.mlp.act | visual.trunk.stages.2.blocks.12.mlp.act | visual.trunk.stages.3.blocks.2.mlp.act | visual.head.mlp.fc2 |
| **zsViT** | visual.transformer.resblocks.0.mlp.gelu | visual.transformer.resblocks.12.mlp.gelu | visual.transformer.resblocks.23.mlp.gelu | visual.ln_post |
| **ftConvNeXt** | stages.0.blocks.0.mlp.act | stages.2.blocks.12.mlp.act | stages.3.blocks.2.mlp.act | head.pre_logits.act |
| **ftViT** | blocks.0.mlp.act | blocks.12.mlp.act | blocks.23.mlp.act | norm |
